# A BLAST from the past: revisiting blastp’s E-value

**DOI:** 10.1101/2024.07.16.603405

**Authors:** Yang Young Lu, William Stafford Noble, Uri Keich

## Abstract

The Basic Local Alignment Search Tool, BLAST, is an indispensable tool for genomic research. BLAST established itself as the canonical tool for sequence similarity search in large part thanks to its meaningful statistical analysis. Specifically, BLAST reports the E-value of each reported alignment, which is defined as the expected number of optimal local alignments that will score at least as high as the observed alignment score, assuming that the query and the database sequences are randomly generated.

Here we critically evaluate the E-values provided by the standard protein BLAST (blastp), showing that they can be at times significantly conservative while at others too liberal. We offer an alternative approach based on generating a small sample from the null distribution of random optimal alignments, and testing whether the observed alignment score is consistent with it. In contrast with blastp, our significance analysis seems valid, in the sense that it did not deliver inflated significance estimates in any of our extensive experiments. Moreover, although our method is slightly conservative, it is often significantly less so than the blastp E-value. Indeed, in cases where blastp’s analysis is valid (i.e., not too liberal), our approach seems to deliver a greater number of correct alignments. One advantage of our approach is that it works with any reasonable choice of substitution matrix and gap penalties, avoiding blastp’s limited options of matrices and penalties. In addition, we can formulate the problem using a canonical family-wise error rate control setup, thereby dispensing with E-values, which can at times be difficult to interpret.

## 1 Introduction

The Basic Local Alignment Search Tool, BLAST, is a cornerstone of genomics research. It enables researchers to search large sequence databases for similar subsequences, facilitating the identification of homologous and orthologous genes, proteins and conserved domains in both inter- and intra-species analyses. As such, BLAST has been instrumental in a wide range of biological research areas, from understanding the evolutionary relationships between species to predicting protein structure and function to designing new drugs. BLAST’s popularity is borne out by the number of citations it has received: the original BLAST paper [3] has over 109,000 citations, and the followup paper [4] has over 84,000 citations (Google Scholar, 1/2024).

BLAST was not the first similarity search tool; however, it quickly became the de facto standard thanks to two key advantages it had over FASTA [19], its key competitor at the time. The first was that BLAST ran a lot faster than FASTA. The second was that BLAST provided the user with meaningful statistical analysis of its output.

BLAST evaluates the significance of a reported local alignment based on the assumption that the distribution of the score of an optimal local alignment, or in BLAST’s terminology, a high-scoring segment pair (HSP), is a Gumbel distribution. This assumption is well grounded in theoretical asymptotic results that cover ungapped alignments [13, 11]. The extension of this significance estimation approach to the more common case of gapped alignments is largely based on empirical evidence [2] and some limited analytic results [5]. In practice, the parameters of the Gumbel distribution must be pre-computed, based on simulations using specific substitution matrix and gap parameters.

The Gumbel-based p-value BLAST computes for an HSP must be adjusted to control for multiple testing: although the p-value provides valuable information about an alignment between the query and a specific database sequence, in a typical application of BLAST we seek to align the query against a database containing many sequences. Accounting for this multiple hypothesis testing scenario is challenging because the database typically contains many closely related sequences. Therefore, standard methods that rely on the independent hypotheses assumption are not valid here.

The solution chosen for BLAST was to express the significance in terms of the expected value, or the *E-value*. Specifically, BLAST’s E-value is defined as the expected number of HSPs that will score at least as high as the observed alignment score, assuming that the query and the database sequences are randomly generated by an independent and identically distributed (iid) process. As explained in Section 2.1 below, BLAST leverages the sequence-specific Gumbel p-value to readily compute the database-wide E-value. This offers BLAST a very simple way to account for all the different possible alignments that the database might offer the query. However, the E-value represents a significant departure from the classical statistical approach to the problem of multiple hypothesis testing, which raises the question of whether one of the more common approaches might still be applicable.

We are now over 30 years past the initial release of BLAST, and in view of the significant advances in computing power we sought to reevaluate the E-value approach. We specifically focused on blastp, the standard protein BLAST that is applicable to amino acid sequences. Via extensive simulated draws from the null we show that, while generally reasonable, blastp’s E-values can at times be overly conservative, while at others, alarmingly, they can be too liberal, i.e., blastp is inflating the significance of the reported alignments. For example, we noted an E-value of 0.05 or smaller reported in over 10% of random alignments.

We therefore offer an alternative significance analysis that relies on generating a sample of size *m* (we used *m* = 50) from the distribution of the maximal alignment score (Figure 1). We then compute a p-value, assessing how unlikely it is that our original maximal alignment score came from the same null sample, assuming that all scores were generated by a Gumbel distribution. This computation is done by a novel adaptation of the rationale behind the independent two-sample t-test to our setting.

**Figure 1:**
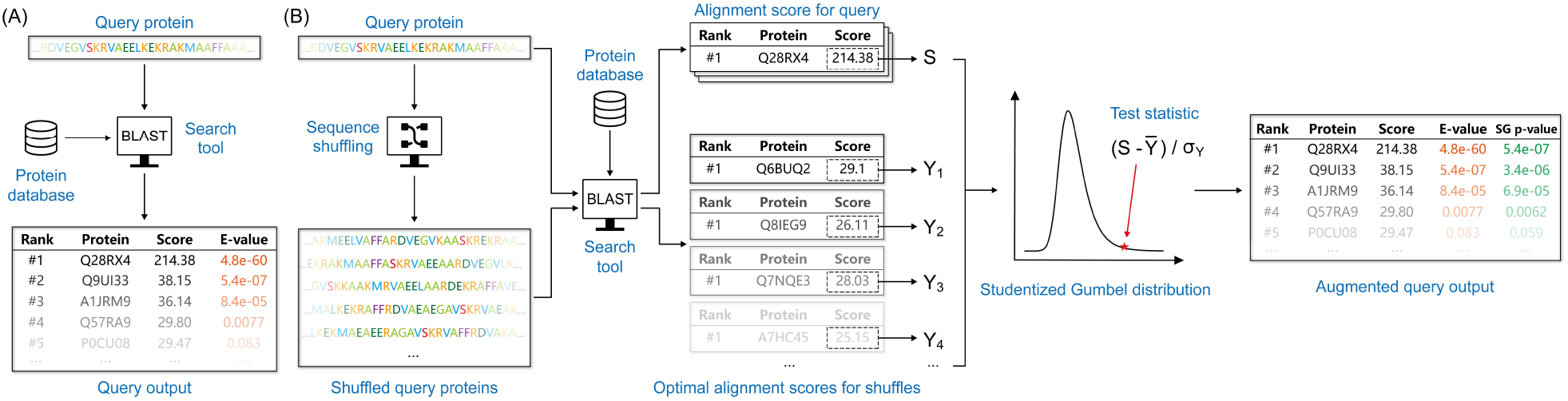
The workflow of (A) blastp and (B) the Studentized Gumbel (SG) p-value calculation.

Compared with blastp’s E-values, our approach has several advantages and one glaring disadvantage. On the positive side, first, our significance analysis is expressed in terms of a classical statistical approach to multiple hypothesis testing: we compute a p-value, and we control the canonical family-wise error rate (FWER). Specifically, if we report all alignments whose p-value is ≤ 0.05, then the probability that even one random alignment to the query is reported is ≤ 0.05. Second, our approach does not rely on any pre-computed parameters, so it is applicable anywhere the null sample generation is feasible. Third, we show that in those cases where blastp’s E-value is statistically valid (i.e., it is not liberal) we deliver more statistical power (i.e., a greater number of interesting alignments). On the negative side our approach imposes a significant runtime penalty (a factor of *m*) and would not have been viable back when BLAST was introduced. However, today, with our vastly improved computing power, the extra computational expense can be easily justified for many applications of blastp.

In addition to comparing with blastp, we include BLAST’s competitor, FASTA, in our analyses. We find that, although FASTA’s E-value is defined and computed differently than BLAST’s, many of the observations stated above hold for FASTA as well. The one exception is that unlike blastp, FASTA also does not rely on pre-computed parameters, and hence its significance analysis is at least as widely applicable as our approach.

For the remainder of this paper when we refer to BLAST we specifically mean blastp. Accordingly, all our analyses were done in the context of databases of sequences of proteins or protein domains. Extending our approach to nucleic acid sequences is a matter for future research.

## 2 Background

### 2.1 Calculating the BLAST E-value

BLAST evaluates the significance of an optimal local alignment (HSP) with score *s* through its E-value, defined as the expected number of HSPs that score ≥ *s*, assuming that the query of length *n*, as well as the *N* database sequences of lengths *l*_1_, *l*_2_, …, *l*_*N*_, are randomly generated. To compute the E-value, BLAST starts with the assumption that the distribution of the score of an HSP is a Gumbel. Specifically, the p-value of an HSP with score *s*, i.e., the probability of finding an HSP with score≥ *s*, in a randomly generated query of length *n* and a *single* database sequence of length *l* can be approximated by

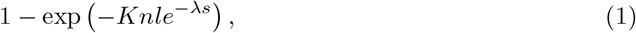

where *K* (related to location) and *λ* (rate, or 1/scale) are the parameters of the Gumbel distribution (precomputed specifically for the considered substitution matrix and gap parameters).

This p-value can next be readily leveraged to compute the expected value of *X*_*s*_, the number of HSPs between such randomly drawn query and database sequence that score≥ *s*. Indeed, it was theoretically established in the ungapped case and empirically observed for the gapped case that *X*_*s*_ has a Poisson distribution [14, 13]. Recalling that for a Poisson(*µ*) random variable *X* we have, *P* (*X >* 0) = 1 − *e*^−*µ*^, and noting that *X*_*s*_ *>* 0 if and only if there exists an HSP that scores ≥ *s*, it follows from (1) that

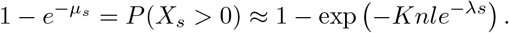

That is, *µ*_*s*_, the expected number of random HSPs between the query and the database sequence that score ≥ *s* is approximated by

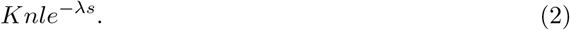

Expectation is additive, so if we add these expectations over all *N* database sequences of lengths *l*_1_, *l*_2_, …, *l*_*N*_, then we get that the expected number of random HSPs between the query and the entire database that score ≥ *s* is approximately

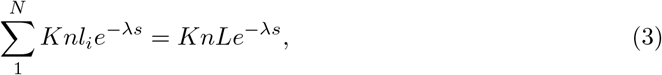

where 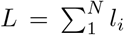 *l*_*i*_ is the total length of the database. This is essentially BLAST’s E-value, except that BLAST employs a sophisticated edge effect correction to account for the fact that an alignment cannot start arbitrarily close to the end of a sequence. Specifically, the *nl* term in (2) is replaced by a complicated term that factors in that edge effect. To compute the final E-value we note that (3) can be derived from (2) by multiplying by *L/l*, so BLAST employs the same adjustment to obtain the overall, database-wide E-value from the edge-corrected, sequence-specific E-value.

### 2.2 Calculating the FASTA E-value

FASTA also reports its significance estimates via the E-value; however, there is a subtle difference between its goal and that of BLAST, which impacts how the two methods evaluate the significance of their respective results. Specifically, BLAST aims to report all sufficiently high-scoring optimal local alignments between the query and the database, and it evaluates each one by computing the expected number of random alignments of the same or higher score. FASTA, on the other hand, aims to find all *database sequences* that are sufficiently similar to the query. “Similar” here means that the two sequences share a sufficiently significant local alignment, and FASTA typically only reports the top alignment between the two sequences. Accordingly, it assesses the significance of the similarity by computing an E-value, which it defines as the expected number of random database sequences that will have the same or higher similarity score.

FASTA also takes a different approach from BLAST in how it computes its E-value. Briefly, FASTA’s default E-value computation (called “REGRESS1”) first regresses the observed similarity scores against the log of the database sequence length, in order to find the mean and variance of the null sequence similarity score as a function of the database sequence length. Here, the underlying assumption is that the vast majority of the database sequences offer a random match to the query. FASTA employs some heuristics to take out of the estimation process the few sequences that might be truly related to the query. The Gumbel distribution is then fitted to the normalized similarity scores, and the fitted distribution is used to estimate the probability of seeing the observed normalized sequence similarity score between the query and a single random database sequence.

To overcome the unknown dependence structure in the database, FASTA converts the single sequence p-value to a database-corrected E-value similarly to BLAST. In FASTA’s case this is done by multiplying the p-value by the number of sequences in the database, which coincides with the Bonferroni correction for multiple testing: one for each database sequence. Importantly, unlike BLAST, FASTA’s significance analysis does not depend on pre-computed parameters, so it can be applied to any combination of a substitution matrix and gap penalties, making it more widely applicable than BLAST.

## 3 Controlling the FWER using Studentized-Gumbel (SG) p-values

In this section we outline our approach, and we provide further details in the subsequent section. Consider a canonical hypothesis testing problem, where you are given an observation *X* and you want to test the null hypothesis that it came from, say, a *N* (*µ, σ*^2^) distribution. If the parameters (*µ, σ*^2^) are known, then you standardize *X* by computing its so-called *Z*-value, *Z* = (*x* − *µ*)*/σ*, and then compute a p-value based on the fact that *Z* ∼ *N* (0, 1).

Suppose next that *µ* and *σ* are unknown but that you can generate a small sample *Y*_1_ …, *Y*_*m*_ from the null distribution. If that null is again a *N* (*µ, σ*^2^) distribution then you can use 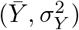, the sample mean and variance of *Y*, in lieu of the unknown (*µ, σ*^2^). Specifically, your test statistic is now the studentized value of *X*, defined as 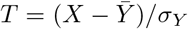. You can next assign a p-value to *X* by realizing that, up to a constant of of 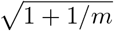 has a t-distribution with *m* − 1 degrees of freedom.^1^

We are interested in testing whether *S, the score of the optimal local alignment across the entire database*, is an observation from the null. Here we define the null as the distribution of the score of the optimal alignment to a random shuffling of the given query. Ideally, we would have searched all possible permutations of the query against the database, noting the optimal match to each of the shuffled queries. This would have allowed us to precisely characterize the null, which is analogous to knowing (*µ, σ*^2^) above.

Of course, this approach is not practical even for very short queries, so instead we opt, as in the unknown (*µ, σ*^2^) case above, to generate a small sample from the null and ask whether *S* and the sample were generated from the same distribution. Specifically, we first generate the null sample by searching each of *m* random shuffles of the query (we used *m* = 50 here) against the database, noting the corresponding scores of the *m* optimal alignments *Y*_1_, …, *Y*_*m*_. We then use the studentized version of 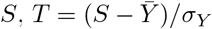, to gauge how well *S* matches the *Y* sample. Specifically, if, as in the normal case, the null distribution of *S* depends only on location and scale/rate parameters, then the distribution of *T* is free of the unknown parameters and hence can be tabulated.^2^

Of course, the Gumbel cumulative distribution function (CDF), *F* (*x*) = exp(*e*^−*λ*(*x*−*µ*)^), is defined only in terms of its rate *λ* and location *µ*, and our empirical analysis below shows that the Gumbel offers a reasonably good fit to the distribution of *S*. Intuitively, this makes sense because BLAST’s entire approach is predicated on the Gumbel approximation to the distribution of the optimal alignment score between two random sequences. Moreover, it is easy to see that a maximum of independent Gumbel random variables (RVs), each with the same rate *λ*, is again a Gumbel RV with the same *λ*.^3^ Therefore, it is reasonable that the distribution of *S*, which is defined by taking the maximum over all database sequences, each with an approximate Gumbel-distributed null optimal score, will again be approximately Gumbel.^4^

In the normal case the studentized 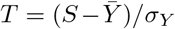 has (up to a constant) the well-tabulated t-distribution with *m*−1 degrees of freedom. As noted, when *S* and *Y*_1_, …, *Y*_*m*_ are Gumbel with location *µ* and a rate *λ*, the distribution of *T* again depends only on *m*; however, to the best of our knowledge, this distribution does not have a name, nor is it tabulated. We therefore refer to it as the “Studentized-Gumbel(*m*)” (SG_*m*_) distribution, and we resorted to using Monte Carlo simulations with importance sampling (detailed below) to estimate its CDF. Figure S3A shows an estimate of the density of SG_50_ using a sample of 10^7^ points.

Having tabulated the SG_50_ distribution, which approximates the null distribution of the studentized version of the database-wide optimal alignment score *S*, we can use it to assign a database-corrected p-value to every BLAST alignment. That is, we can estimate the probability that a random shuffle of the query will have an alignment scoring as least as high as observed. As we show below, if we then report all the alignments whose p-value is ≤ *α* then we control the FWER among the reported alignments at that level, i.e., the probability that even one random alignment is reported is ≤ *α*.

## 4 Methods

### 4.1 Computing the Studentized Gumbel (SG) distribution

We used Monte Carlo simulations with importance sampling to estimate the CDF of SG_50_ as follows. We divided the range of positive values *T* can attain between 0 and 45 into bins of size 0.001. The upper limit of 45 was arbitrarily chosen (more on that below), and the lower limit of 0 was chosen so as not to waste computational resources on negative values of *T*: if your best score is less than the mean of a null sample then it is probably not an interesting one.

We next drew *N* = 10^10^ samples of size *m* = 50 from the Gumbel distribution with *µ* = 0 and scale 1*/λ* = 3 (we could have used any *λ*, except this choice impacts the parameters used in the importance sampling step). This gave us *N* independent draws (*µ*_1_, *σ*_1_), …, (*µ*_*N*_, *σ*_*N*_) of the sample mean 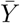, and the sample standard deviation *σ*_*Y*_ of the null samples of *Y*_1_, …, *Y*_*m*_. If we had drawn the values of *S* from the same Gumbel distribution, then we would have been rather limited in our ability to compute very small p-values (corresponding to very large values of *S*), so instead we used importance sampling, as explained next.

We drew *N* values of *S* from a Gumbel distribution with *µ* = 33 and scale 1*/λ* = 15. This significant shift in location allowed us to sample much higher values of *S* than had we used the same *µ* = 0 that was used to generate the *Y* sample. At the same time we increased the scale (the reciprocal of the rate), so as to make sure we are effectively sampling *S* across a wide range of values. To account for the fact that *S* was not sampled from a Gumbel(0, 1*/*3), we weighted each observed value *s*_*i*_ by the log-likelihood ratio (aka the Radon-Nikodym derivative), *f*_*G*(0,1*/*3)_(*s*)*/f*_*G*(33,1*/*15)_(*s*), where *f*_*G*(*µ,λ*)_(*s*) = *λ* exp[−*λ*(*s* − *µ*) − *e*^−*λ*(*s*−*µ*)^] is the Gumbel(*µ, λ*) density. That weight was assigned to the bin that contained the studentized value of *s*_*i*_: *t*_*i*_ = (*s*_*i*_ − *µ*_*i*_)*/σ*_*i*_.

For each bin we then considered the weights *w*_1_, …, *w*_*n*_ of all the studentized samples that fell in that bin, or in bins corresponding to higher studentized values. If we let *t* denote the center of the considered bin, then the right tail of the CDF at *t* is estimated as 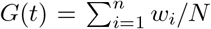. Additionally, we computed the sample standard deviation of those same weights, and the first bin for which the ratio of this standard deviation to *G*(*t*), equivalently the coefficient of variation, was ≤ 0.01 defined our precision cutoff. That is, we did not try to estimate the tail probability, *G*(*t*) for values of *t* greater than that cutoff. In practice we further truncated the cutoff of 34.599 to 34. Figure S3B shows the (log10) of the computed tail probabilities.

### 4.2 Computing the SG_*m*_ p-values

Equipped with the right tail probabilities of the SG_*m*_ distribution it is conceptually straightforward to compute the p-values of the observed scores *S*_1_, …, *S*_*j*_ of aligning a given query to the database. Specifically, we shuffle the same query *m* times (here we use *m* = 50) and apply BLAST (or more generally, the search tool) to the *m* shuffled queries in exactly the same way it was applied to the original query.

Let *Y*_1_, …, *Y*_*m*_ denote the *m* scores of the maximal alignments for each of the *m* shuffles. We use the sample moments of *Y* to studentize the observed scores and then look up the corresponding entries in the table of SG_*m*_ tail probabilities that we computed in the previous section. Note that any studentized value that exceeds the accuracy cutoff of 34 is assigned a p-value of *G*(34)=8.716e-15. Similarly, any value ≤ 0 is assigned a p-value of 1. An algorithmic description of this process is available in Algorithm 1.

### 4.3 Controlling the family-wise error rate (FWER)

We first describe a straightforward procedure that determines which of the *k* alignments of the given query to the database that BLAST found should be reported as significant. Let *S*_1_, …, *S*_*k*_ be the corresponding scores of those local alignments returned by BLAST, and let *α* be the selected significance threshold (canonically *α* = 0.05). Then,

1. Apply Algorithm 1 to obtain the SG_*m*_ p-values *p*_1_, …, *p*_*k*_ of the corresponding alignment scores *S*_1_, …, *S*_*k*_.
2. Report as significant any alignment with p-value *p*_*i*_ ≤ *α*.

≤

We next argue that, if we assume that our SG p-values are valid, that is, that under the null hypothesis *P* [p-value(*S*) ≤ *α*] ≤ *α*, then our procedure controls the FWER among the reported local alignments to the given query: the probability that even one random/null-generated alignment is reported is ≤*α*.

Note first that because we are only concerned about reporting random alignments, we can assume without loss of generality the worst case scenario that all alignments to the query are null generated, or equivalently, that the query is randomly shuffled. Let 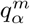denote the 1 – *α* quantile of our SG_*m*_ distribution, *S* = max *S*_*i*_, and 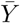 and *σ*_*Y*_ denote the sample moments of the randomly drawn *Y*_1_, …, *Y*_*m*_ in Algorithm 1. Then

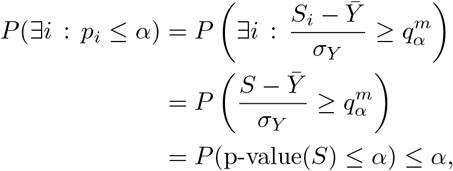

where the last inequality follows by our assumption that the SG p-values are valid. We conclude this section with two comments:

1. The above discussion is predicated on our SG p-values being valid. First note that, ignoring the sampling errors when estimating the SG_*m*_ distribution (Section 4.1), if the null distribution of the maximal alignment score *S* is indeed a Gumbel then our p-values are valid. Our simulated draws show that, just as in the case of BLAST, this is a reasonable assumption for longer queries (Section 5.2). Moreover, even when the fit to the Gumbel is not perfect, which is evident in some cases with shorter queries, in practice our SG p-values are conservatively biased (Section 5.3), and hence, still valid even in those cases.
2. While we mostly referred to BLAST throughout this section, its content applies just as well to other alignment/similarity search tools including FASTA, SSEARCH (A SmithWaterman alignment algorithm implemented in the FASTA program package [18]) and AB-BLAST (a rebranded version of WU-BLAST), as long as the SG p-values are valid — a point which we address empirically below.

### 4.4 Comparing the SG_50_ p-values with the E-values

To compare our SG p-values with E-values we used two types of experiments. The first was designed to test the validity of the reported values; the second compares their statistical power, asking how effectively they can help us reject the null when we should.

#### 4.4.1 Analyzing the validity of the significance measures

We empirically studied the validity of the SG_50_ p-values and the reported E-values by simulating draws from the null distribution of optimal alignments by searching randomly shuffled sequences against a database. Specifically, we used multiple setups, where we varied the search engine, the selected database, the query that was shuffled, the substitution matrix and the gap penalties. For each such setup we generated a sample of *N*_*s*_ = 10^6^ draws from the null distribution by applying the search tool with the chosen parameters to report the maximal alignment score, *S*, as well as the minimal alignment E-value, *E*, for each of the *N*_*s*_ shuffled sequences.^5^ The following options were considered for each category when defining a setup.

- Search engine: NCBI BLAST (versions 2.11.0 and 2.15.0), AB-BLAST (version 3.0, which we ran with the `-kap` option because its default sum statistic is not compatible with our p-value approach), FASTA, and SSEARCH (both version 36). The tools were run using their default settings except where explicitly stated otherwise.
- Database: the human-annotated SCOP database [16] (release 2022-06-29, consisting of 35,644 family-level representative domain sequences), the Swiss-Prot database [6] (release 2023_01, consisting of 481,450 manually annotated and non-redundant protein sequences), and the ASTRAL40 database [9] (version 2.08, a subset of the SCOP database containing 15,178 domain sequences, each with less than 40% identity to the others).
- Query^6^: one of two sets of five queries of varying lengths. The first set consisted of five SCOP domains that were randomly selected subject to having lengths of 45, 90, 175, 350 and 700 amino acids (Table S1). The UniProt IDs of the proteins containing those five domains are: Q88D80, Q9NSN8, Q9D8T0, P17654 and A8FDC4. The second set was made of five sequences of the same lengths as of the first set but each sequence was randomly drawn by an iid process using the marginal amino acid frequency table that is specified in the file `blast_stats.c` of BLAST’s source code (Table S2). When we refer to a query by its length we mean one from the first set; we will explicitly mention “iid” in the few cases where we refer to the second set.
- Substitution matrices: BLOSUM45 (B45), BLOSUM50, BLOSUM62, BLOSUM80, and BLOSUM90 of [12], PAM30, PAM70, and PAM250 of [10], and the non-standard PFA-SUM60 of [15].
- Gap penalties: largely the default for the chosen substitution matrix, e.g., BLAST’s default for BLOSUM62 is (11,1): 11 to open a gap and 1 to extend it. For PFASUM60 we used the (15,1) penalty recommended by its authors [15]

We next used these null samples to graphically examine how well does the Gumbel distribution model the null distribution of the optimal score, as well as the validity of the SG_50_ p-values and the E-values. Specifically, we used probability plots specifying, for each of the *N*_*s*_ shuffled queries, the (log10 of the) fraction of the shuffles (x-axis) for which the optimal alignment p-value/E-value is better than the (log10 of the) the optimal p-value/E-value associated with that shuffled query (y-axis).

We organized the probability plots in 3-column wide figures of plots as follows:

- Each row is dedicated to a different shuffled query, and the rows are ordered in increasing query length. Note that the queries of length 700 were left out of the figures due to space constraints.
- The left, “Gumbel p-values” column examines how well the null distribution of *S* fits the Gumbel distribution by looking at the frequency of p-values, computed using the Gumbel distribution with parameters estimated via maximum-likelihood estimation (MLE) from the entire sample of *N*_*s*_ points. Note that the closer the points are to the diagonal line the better the fit is, and that Figure S2A provides a reference point to what can be expected when the fit is perfect.
- The middle, “SG p-values” column analyzes *N*_*s*_ SG_50_ p-values, one computed for each optimal alignment score *S*. In this case, the auxiliary null samples needed for studentizing each observed *S* were randomly drawn with replacement from the set of *N*_*s*_ optimal scores. A curve that goes above the diagonal indicates conservative, and hence valid p-values, although if it is significantly above the diagonal its power might be reduced. On the other hand, if the curve dips significantly below the line, then the p-values are most likely invalid.
- The right, “E-values” column examines the frequency of the reported minimal E-values. Because an E-value should be greater than its associated p-value, an E-value of, say 0.01 or less, should not appear in significantly more than 1% of the null samples. Thus, dips below the diagonal in the right column panels indicate a potential liberal bias: the reported E-values unduly inflate the significance of the alignments. A conservative bias is more difficult to quantify because the E-value is an overestimate of a p-value, however points substantially above the line indicate potential conservative bias which would translate to reduced power.

For example, the bottom row of Figure 2 is based on data generated by using BLAST to search 10^6^ shuffles of a query of length 175 amino acids against the Swiss-Prot database [6], with PAM70 and its BLAST default gap penalties of (10,1). The top row was derived using a similar sample of size 10^6^ from the null, but this time BLAST was used to search the same shuffled queries against the SCOP database [16] with BLOSUM45 and its BLAST default gap penalties of (14,2).

**Figure 2:**
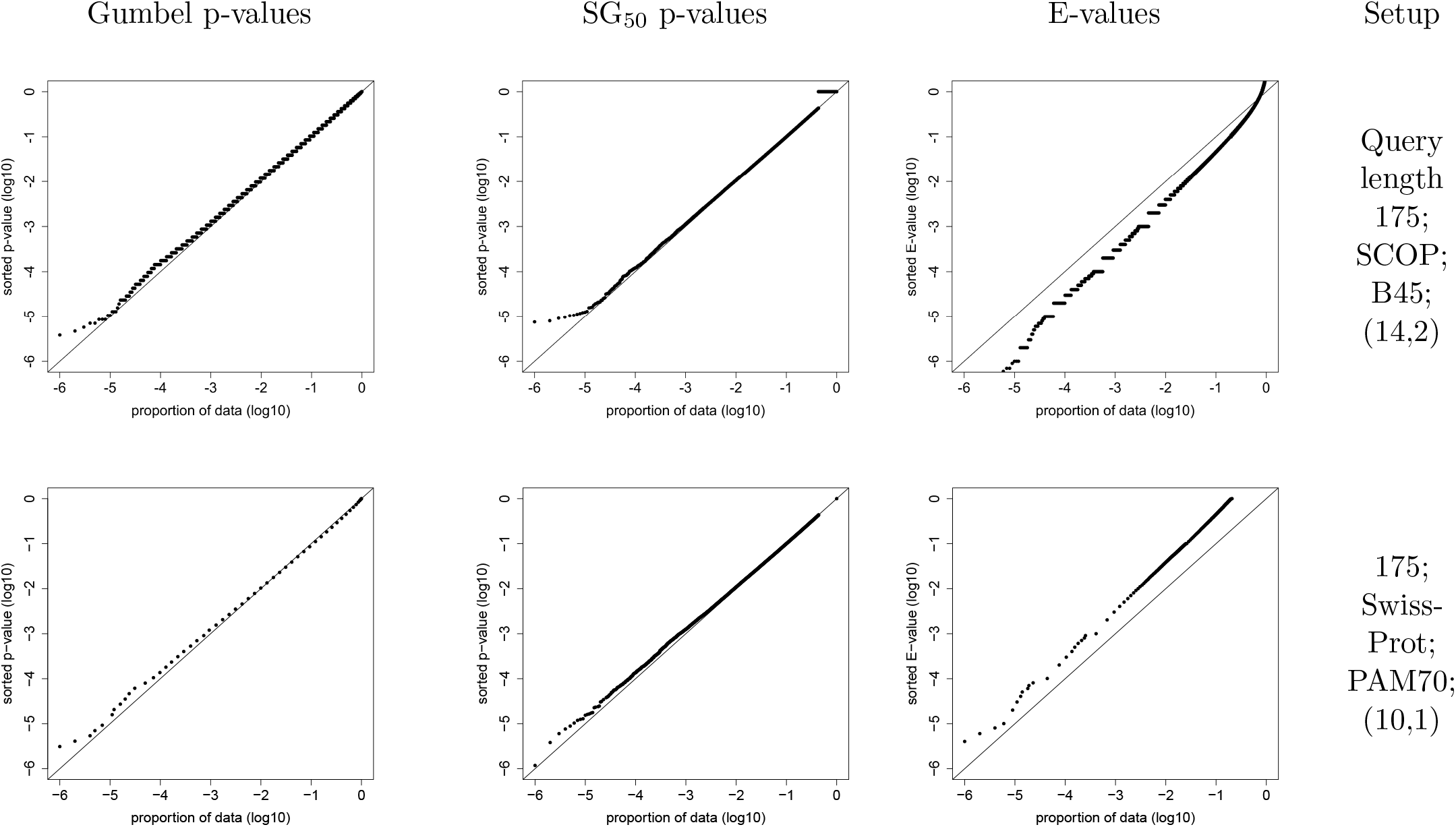
BLAST E-value can be liberal as well as conservative. The top row plots were made based on a sample generated by using BLAST to search 10^6^ shuffles of a query of length 175 amino acids against the SCOP database with BLOSUM45 and its BLAST default gap penalties of (14,2). The bottom row is similarly based on 10^6^ BLAST searches against the Swiss-Prot database with PAM70 and its BLAST default gap penalties of (10,1). The left panel examines the fit of the optimal alignment score to the Gumbel, the middle one the validity of our SG_50_ p-values, and the right panel examines the validity of the E-values (see Section 4.4.1 for details).

In addition, for the canonical significance cutoffs of *α* ∈ *{*0.05, 0.01, 10^−3^, 10^−4^, 10^−5^, 10^−6^*}* we looked at the frequency with which the SG_50_ p-values / E-values are ≤ *α*. Specifically, if that frequency is *> α*, then we conduct a 1-sided binomial test to see if this liberal tendency is statistically significant in which case it is explicitly mentioned, although this occurs too commonly for E-values to report every such violation.

#### 4.4.2 Comparing the power of the E-values and SG_50_ p-values

To compare the statistical power of the significance measures we performed homology search experiments based on the ASTRAL40 database [9] (version 2.08). ASTRAL40 is widely recommended as the gold standard for evaluating homology search performance [8, 20].

The domains in the ASTRAL40 database are hierarchically organized into classes, folds, superfamilies, and families. To assess the sensitivity of the homology search in this experiment we made the common assumption that two domains from the same superfamily are homologous (positive).

We performed a homology search for each sequence within the ASTRAL40 database against the entire database, employing the same versions of BLAST, AB-BLAST, FASTA and SSEARCH noted in Section 4.4.1. AB-BLAST, FASTA were applied using BLOSUM62 (11,1), PAM70 (10,1), and the non-standard PFASUM60 (15,1) BLAST cannot be used with non-standard matrices so we only used the first two matrices with it. SSEARCH was used with PFASUM60 (15,1).

Power was compared in terms of the number of positive alignments that were reported at the cutoffs of 0.01 and 0.05. Based on our validation experiment reported below, we did not include AB-BLAST’s E-values in this analysis because they are clearly invalid (Figure S29).

## 5 Results

### 5.1 The E-values can be too liberal

The right columns of Figure 2 and Figures S5-S35 provide evidence that all the different flavors of E-values can be too conservative in some setups and too liberal in others. To highlight a few of the results, starting with BLAST, we see in the top right column of Figure 2 that BLAST can substantially inflate the significance of the alignments. For example, 11% of the optimal alignments (obtained by running BLAST on shuffles of the length 175 query against the SCOP database using BLOSUM45 (14,2)) have an E-value ≤ 0.05 (a p-value of 0 according to a 1-sided binomial test), and 2.7% have an E-value ≤ 0.01 (a p-value of 0). The bottom right panel of that figure shows the flip side where, for example, only 1.3% of the optimal alignments (generated by running BLAST on shuffles of the same length 175 query against the Swiss-Prot database using PAM70 (10,1)) have an E-value ≤ 0.5. Moreover, Figures S5-S7 indicate that the E-values often substantially inflate the significance of optimal alignments found using BLOSUM45 (14,2), and Figures S26-S28 indicate that the same applies to alignments generated with PAM250 (14,2). Similarly, while not as severe, Figure S12 shows cases of over- and under-estimation of the E-value even using the default BLOSUM62 (11,1).

Moving on to AB-BLAST we were surprised to see that its reported E-values are clearly problematic. For example, Figure S29 shows that searching shuffles of the length 45 query against the SCOP database using BLOSUM62 (9,2)^7^ the reported E-values are too liberal for moderately small E-values (e.g., 6.1% of the alignments have an E-value ≤ 0.05; p-value of 0), but they become very conservative for smaller E-values (e.g., none of the 10^6^ alignments has an E-value ≤ 0.0001). The situation is even worse for shuffles of the length 90 query where, for example, 35% of the alignments have an E-value ≤ 0.05 (p-value of 0). Note that while these results were obtained using the `-kap` option, we saw similar problems when using AB-BLAST’s default sum statistic. For this reason we chose to exclude AB-BLAST’s E-values when comparing the power of the significance measures.

FASTA and SSEARCH rely on the same method when computing their E-values. While generally they tend to be overly conservative, we found examples of both tools inflating the significance. Indeed, the right column of Figure S35 shows that SSEARCH (using PFASUM61 (15,1) and the ASTRAL40 database) can be very conservative for a query of length 45 (e.g., only 3 of the 10^6^ alignments have an E-value ≤ 0.001), while for a query of length 350 it is too liberal (e.g., 6.5% of the alignment have an E-value ≤ 0.05; p-value of 0). A similar trend, though less pronounced, can be observed for FASTA in Figure S32, where, for example, for shuffles of the length 350 query (searched against ASTRAL40 using BLOSUM62 (11,1)), 6% of the alignments have an E-value ≤ 0.05 (p-value of 0).

In addition to its default procedure for computing its E-value, FASTA also offers shuffled-based procedures. We looked at two of them, `-z 11` and `-z 21`, which are the analogs of the default procedure applied to shuffled database sequences rather than the original sequences. The two methods differ in their selection of sequences that are shuffled. We found that in both cases the resulting E-values can be too liberal, particularly so for the `-z 21` option, e.g., using FASTA to search 10^6^ shuffles of the length 700 query against the ASTRAL40 database with the non-standard PFASUM60 (15,1) we find 6.1% of the `-z 11` generated E-values and 36% of the `-z 21` generated E-values are ≤ 0.05 (both binomial test p-values are 0).

### 5.2 The optimal alignment score can be reasonably modeled by the Gumbel distribution

The left column of Figure 2 shows the fits of the optimal alignment score, *S*, to the Gumbel distribution (based on 10^6^ samples) for a couple of examples that we highlight.

Additionally, the left columns of Figures S5-S35 demonstrate that, consistent with our expectation, the Gumbel fit generally improves with increasing query length. Keeping in mind the stochastic nature of the sample, we provide in Figure S2A an example of what those probability plots should look like when the sample of 10^6^ points is indeed generated from the Gumbel distribution. With that reference figure in mind, it is clear that the fit for the length 45 shuffles is often far from perfect — a fact we will return to below.

### 5.3 The SG_50_ p-values appear to be valid

As noted above, the Gumbel distribution is not an ideal model for the null distribution of *S* for shorter query sequences. Fortunately, as can be verified by the middle panel of the top row of Figures S5-S35, this results in a consistent conservative bias, which means the SG_50_ p-values are still valid. Moreover, while these p-values of the length 45 shuffles are conservative, they are still typically significantly less so compared with the corresponding E-values. More generally, the SG_50_ p-values are much more consistent than the E-values in the same experimental setup. For the longer queries, together with the improved fit between the Gumbel and the null distribution of the maximal alignment score, the SG_50_ p-values improve considerably, while still remaining valid. On the latter point we note that, in contrast with the E-values, across the dozens of experimental setups described in S5-S35 the fraction of p-values that were ≤ *α* rarely exceeded *α* (for *α* ∈ { 0.05, 0.01, 10^−3^, 10^−4^, 10^−5^, 10^−6^}). Moreover, out of those only once was the 1-sided binomial test significance below the standard 0.05 cutoff: 0.0487. After accounting for the hundreds of tests involved (6 level of *α* for each of the dozens of setups), this can be safely ruled as statistically insignificant.

### 5.4 The SG_50_ p-values discover more homologous sequences when the E-values not too liberal

The SG_50_ p-values rely on an auxiliary null sample, which could have theoretically compromised their statistical power. Thus, it is particularly reassuring to see that in our power comparisons they typically detect more homologous sequences than the E-values do. Specifically, examining Table 1 we find that the number of reported homologs using the p-value cutoff is typically larger than the corresponding number reported using the E-value cutoff. For example, using SSEARCH and PFASUM60 (15,1) we report 145,288 homologs using the E-value cutoff of 0.01 compared with 149,736 using the p-value cutoff of 0.01 (a 3% increase). The exception is when using BLAST with BLOSUM62 (11,1), where it reports more homologs using the E-value cutoff criterion (140,488 vs. 136,921 or 2.6% more). However, looking at Figure S30 it is also clear that BLAST inflates the significance of some of its alignments in this setup (e.g., for a query length of 175, 6.1% of the null alignments have an E-value ≤ 0.05; a p-value of 0), so any advantage the E-value has in this setup should be taken with a grain of salt.

**Table 1:** The number of ASTRAL40 homologous sequences reported at the given threshold. See Section 4.4.2 for details.

| Search tool | Substitution matrix<br>(Gap open/extension penalty) | The number of reported homologs |  |  |  |
| --- | --- | --- | --- | --- | --- |
|  |  | E-value <0.01 | SG<sub>50</sub> p-value <0.01 | E-value <0.05 | SG<sub>50</sub> p-value <0.05 |
| AB-BLAST | BLOSUM62 (11,1) | N/A | 138118 | N/A | 151914 |
| BLAST | BLOSUM62 (11,1) | 140488 | 136921 | 149846 | 147360 |
| FASTA | BLOSUM62 (11,1) | 128174 | 133185 | 139004 | 144120 |
| AB-BLAST | PAM70 (10,1) | N/A | 95183 | N/A | 104378 |
| BLAST | PAM70 (10,1) | 113522 | 116094 | 119980 | 123767 |
| FASTA | PAM70 (10,1) | 88706 | 91648 | 97284 | 100848 |
| AB-BLAST | PFASUM60 (15,1) | N/A | 146834 | N/A | 159705 |
| FASTA | PFASUM60 (15,1) | 133351 | 137729 | 144578 | 148840 |
| SSEARCH | PFASUM60 (15,1) | 145288 | 149736 | 157231 | 162083 |

Notably, using SSEARCH with PFASUM60 (15,1) and the SG_50_ p-value cutoffs delivers the largest number of discoveries overall, demonstrating the utility of our approach that (a) extends to non-standard matrices, and (b) delivers powerful and valid significance analysis. On the latter point we note that Figure S35 graphically demonstrates the validity of our p-values in this specific setup.

## 6 Discussion

E-values have allowed BLAST to present scientists with a meaningful statistical evaluation of reported local alignments. In this paper we exposed some deficiencies in how the E-values are computed in practice. Specifically, we found that at times they are under-estimated, inflating the significance of the reported alignment, while at others they seemed to be over-estimated. We showed that these problems are shared with other commonly used similarity tools that rely on E-values.

Examining the right (E-values) column of our 3-column figures (e.g., Figure S5) casts doubts on BLAST’s paradigm of computing the E-value based on precomputed Gumbel parameters for the given substitution matrix and gap penalties. Granted, BLAST employs sophisticated methods developed to adjust for the query length (by applying an edge effect correction, e.g., [1]), and for the query compositional bias (e.g., [4]); however, the length adjustment in particular seems to fall short (e.g., Figure S7). In addition, comparing the E-values column of Figures S5 and S6 suggests a database effect that is not properly controlled for.

Our approach tries to address those issues by first observing that the null Gumbel distributional assumption can be reasonably extended from the sequence to the database level. This allows us to compute a p-value in lieu of the E-value. Then, by studentizing the score we get around the problem that the parameters of the said Gumbel distribution vary with the query, as well as with the alignment parameters. As an aside, note that as an alternative to studentizing the observed score *S*, we could also use our auxiliary null sample to estimate the Gumbel parameters and then use the estimated Gumbel CDF to compute a p-value. The problem with this approach is that these “MLE p-values” are not valid, as can be verified in Figure S4.

Aside from the issue of whether or not the E-values are computed accurately enough, we question whether this is the right notion of significance for this context. One of the arguments that NCBI makes in favor of using the E-values is that “it is easier to understand the difference between, for example, E-value of 5 and 10 than P-values of 0.993 and 0.99995” https://www.ncbi.nlm.nih.gov/BLAST/tutorial/Altschul-1.html. However, this begs the question of whether one should be interested in alignments that score so low that a random alignment has a probability of 0.993 to score higher.

Moreover, suppose that you found 40 alignments with an E-value of 10. In this case you would be tempted to think this is somehow significant, because you expected only 10 such alignments but you found 40. However, assigning significance to this 40 vs. 10 is a different question and might be simply a reflection of the redundancy in the database.

Instead, we promote here the use of the canonical approach of FWER control based on p-values that are adjusted to the multiple hypothesis problem at hand. Our p-values are empirically valid and are exact in the limiting Gumbel distribution case, while being more conservative with shorter query lengths. Still, we show that they deliver more power than when using E-values, in the case that the latter are not overly liberal.

Properly controlling the FWER when simultaneously searching many queries against one or more databases will typically be too restrictive. Indeed, the common approach to the general multiple testing problem is to switch from controlling the FWER to controlling the false discovery rate (FDR) when facing a large number of hypotheses. Similarly, in [17] the authors suggest controlling the local FDR instead of E-values in a related context of stratified protein domain prediction. Notably, our approach allows us to readily implement FDR control by using our generated p-values as input to a procedure that controls the FDR for dependent hypotheses as well [7].

Our approach is flexible in terms of the substitution matrix and gap penalties it allows, although it is clear that unreasonable penalties or matrices could break the underlying Gumbel approximation. Finally, while it comes with a hefty computational penalty, with a runtime penalty factor of 50 there are still many use cases where our approach is practically applicable. This is in contrast to related works that used sophisticated but impractical Monte Carlo methods to more accurately assess the significance of optimal local alignments, e.g., [21].

To assist users with applying our approach, we developed a wrapper script that first invokes blastp and then runs Algorithm 1 to compute the SG_*m*_ p-values for each alignment that BLAST reports. The interface of the wrapper script, as well as a sample output is available in Section S1.2, while the Apache licensed wrapper itself, as well as a demo are available at https://github.com/batmen-lab/SGPvalue.

## Footnotes

1 One way to see this is to note that 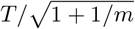 coincides with the test statistic of the independent twosample t-test comparing the *X* and *Y* samples (of sizes 1 and *m*), and the latter statistic has a t-distribution with *m* + 1 − 2 = *m* − 1 degrees of freedom.

2 Indeed, the same location is shared between *S, Y*_*i*_ and 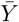; hence, it cancels out in 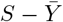and in *σ*_*Y*_. Similarly, the scale is shared between *S, Y*_*i*_, 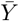 and *σ*_*Y*_; hence, it cancels out in *T*.

3 *P* 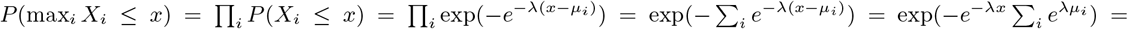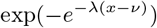 where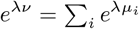, or 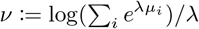.

4 Keepin mind that BLAST assumes that *λ* depends only on the substitution matrix and the gap penalties so it should be shared among all database sequences.

5 Due to computational constraints, in a few setups we used only *N*_*s*_ = 2 10^5^ shuffles.

6 Each of the ten queries had a single set of 10^6^ random shuffles that was shared among all setups that used shuffles of that query

7 We used AB-BLAST’s default gap penalties here.

