## Supplement for "A BLAST from the past: revisiting blastp’s E-value"

#### S1 Supplementary Material

#### S1.1 The algorithm to compute the SG$_m$ p-values

**Algorithm 1:** The SG p-value algorithm

```
Data: A query protein q, a protein database D, a search algorithm S(q, D) that returns a ranked list
      of alignment scores, a list of observed alignment scores s = (s_1, ..., s_k), a specified number of
      shuffles m, T_p the two-column table of right tail probabilities of SG_m computed in Section 4.1:
      first column is the bin center, the second is the tail probability.
Result: PV = (p_1, ..., p_k) - the list of SG_m p-values of (s_1, ..., s_k)
Y ← [];
for i = 0; i < m; i++ do
    q̃ ← shuffle(q);
    Add Y_i = max (S(q̃, D)) to Y;
end
PV ← [];
Calculate the mean Ȳ and the standard deviation σ_Y of Y;
for j = 0; j < k; j++ do
    Calculate the test statistic t_j = (s_j − Ȳ)/σ_Y;
    if t_j ≤ 0 then
        p_j = 1;
    end
    else if t_j > m_x := max T_p[, 1] then
        /* m_x is the maximal bin recorded in the SG_m table T_p                              */
        p_j = T_p[m_x, 2];
        /* The minimum p-value recorded in T_p                                                */
    end
    else
        Find the bin i_j in which t_j falls (i.e., T_p[i_j, 1] ≤ t_j ≤ T_p[i_j + 1, 1]) and set p_j = T_p[i_j, 2];
    end
    Add p_j to PV;
end
```

#### S1.2 The `blastp` wrapper

We implemented Algorithm 1 as part of a wrapper script for `blastp`. Essentially, users can seamlessly invoke `blastp` through the wrapper, resulting in a more reliable statistical estimation in terms of SG-pvalue (Figure S1). The Apache licensed source code is available at (`https://github.com/batmen-lab/SGPvalue`).

**Usage**:
python blastp_wrapper.py [sgpvalue_options] blastp [blastp_options]

**Options**:

**sgpvalue_options**:

| | |
|---|---|
| –num_shuffle <NUM_SHUFFLE> | The number of null |
| –lookup_url <LOOKUP_URL> | The SG look-up table |
| –blastp_url <BLASTP_URL> | The path to the blastp executable file |

**blastp_options**:

| | |
|---|---|
| -db <DB> | BLAST database name |
| -query <QUERY> | Query file name |
| -out <OUT> | Output file name |
| -outfmt <OUTFMT> | Alignment view options |
| -matrix <MATRIX> | Scoring matrix name |
| -evalue <EVALUE> | Expect value (E) for saving hits |
| -num_descriptions <NUM_DESCRIPTIONS> | Show descriptions for this number of database sequences |
| -num_alignments <NUM_ALIGNMENTS> | Show alignments for this number of database sequences |
| -max_target_seqs <MAX_TARGET_SEQS> | Number of aligned sequences to keep |
| -max_hsps <MAX_HSPS> | Maximum number of HSPs (alignments) to keep |
| -word_size <WORD_SIZE> | Word size of initial match. Valid word sizes are 2-7 |
| -gapopen <GAPOPEN> | Cost to open a gap |
| -gapextend <GAPEXTEND> | Cost to extend a gap |
| -threshold <THRESHOLD> | Minimum score to add a word to the BLAST lookup table |
| -comp_based_stats <COMP_BASED_STATS> | Use composition-based statistics |
| -xdrop_gap_final <XDROP_GAP_FINAL> | Heuristic value (in bits) for final gapped alignment |
| -window_size <WINDOW_SIZE> | Multiple hits window size, use 0 to specify 1-hit algorithm |

```
BLASTP 2.14.0+


Reference: Stephen F. Altschul, Thomas L. Madden, Alejandro A.
Schaffer, Jinghui Zhang, Zheng Zhang, Webb Miller, and David J.
Lipman (1997), "Gapped BLAST and PSI-BLAST: a new generation of
protein database search programs", Nucleic Acids Res. 25:3389-3402.


Reference for composition-based statistics: Alejandro A. Schaffer,
L. Aravind, Thomas L. Madden, Sergei Shavirin, John L. Spouge, Yuri
I. Wolf, Eugene V. Koonin, and Stephen F. Altschul (2001),
"Improving the accuracy of PSI-BLAST protein database searches with
composition-based statistics and other refinements", Nucleic Acids
Res. 29:2994-3005.



Database: scop_fa_represeq_lib_latest_all.fasta
           35,644 sequences; 6,930,552 total letters



Query= 8107723_t3_origin

Length=276
                                                                      Score     E
Sequences producing significant alignments:                          (Bits)  Value

8107723 FA=4002088 FA-PDBID=7OIT_AAA FA-UNIID=A0A140TAH5              34.7    0.034
8093645 FA=4002088 FA-PDBID=6QH6_M FA-UNIID=Q3ZC13                    34.3    0.035
8023630 FA=4002088 FA-PDBID=2VGL_M FA-UNIID=P84092                    34.3    0.036
8023950 FA=4002088 FA-PDBID=1H6E_A FA-UNIID=Q96CW1                    33.9    0.051
8021938 FA=4000912 FA-PDBID=1EL0_A FA-UNIID=P22362                    29.1    0.39
8026973 FA=4003124 FA-PDBID=1PYF_A FA-UNIID=P46336                    30.7    0.56
8055168 FA=4007555 FA-PDBID=1WJ3_A FA-UNIID=Q9P232                    28.3    1.5
8099493 FA=4000029 FA-PDBID=6ZZP_A FA-UNIID=Q4FRT2                    28.7    2.3
8082470 FA=4000945 FA-PDBID=4V6U_BM FA-UNIID=Q8U2F9                   28.3    2.5
8051630 FA=4007439 FA-PDBID=5U0P_Q FA-UNIID=P87306                    28.3    3.5
8095272 FA=4003216 FA-PDBID=6OVI_A FA-UNIID=Q5ZYF2                    27.9    3.8
```

**A:** blastp

```
BLASTP 2.14.0+


Reference: Stephen F. Altschul, Thomas L. Madden, Alejandro A.
Schaffer, Jinghui Zhang, Zheng Zhang, Webb Miller, and David J.
Lipman (1997), "Gapped BLAST and PSI-BLAST: a new generation of
protein database search programs", Nucleic Acids Res. 25:3389-3402.


Reference for composition-based statistics: Alejandro A. Schaffer,
L. Aravind, Thomas L. Madden, Sergei Shavirin, John L. Spouge, Yuri
I. Wolf, Eugene V. Koonin, and Stephen F. Altschul (2001),
"Improving the accuracy of PSI-BLAST protein database searches with
composition-based statistics and other refinements", Nucleic Acids
Res. 29:2994-3005.



Database: scop_fa_represeq_lib_latest_all.fasta
           35,644 sequences; 6,930,552 total letters



Query= 8107723_t3_origin

Length=276
                                                                      Score     E    SG
Sequences producing significant alignments:                          (Bits)  Value p-value

8107723 FA=4002088 FA-PDBID=7OIT_AAA FA-UNIID=A0A140TAH5              34.7    0.034 0.058
8093645 FA=4002088 FA-PDBID=6QH6_M FA-UNIID=Q3ZC13                    34.3    0.035 0.071
8023630 FA=4002088 FA-PDBID=2VGL_M FA-UNIID=P84092                    34.3    0.036 0.071
8023950 FA=4002088 FA-PDBID=1H6E_A FA-UNIID=Q96CW1                    33.9    0.051 0.087
8021938 FA=4000912 FA-PDBID=1EL0_A FA-UNIID=P22362                    29.1    0.39  0.432
8026973 FA=4003124 FA-PDBID=1PYF_A FA-UNIID=P46336                    30.7    0.56  0.417
8055168 FA=4007555 FA-PDBID=1WJ3_A FA-UNIID=Q9P232                    28.3    1.5   0.432
8099493 FA=4000029 FA-PDBID=6ZZP_A FA-UNIID=Q4FRT2                    28.7    2.3   0.432
8082470 FA=4000945 FA-PDBID=4V6U_BM FA-UNIID=Q8U2F9                   28.3    2.5   0.432
8051630 FA=4007439 FA-PDBID=5U0P_Q FA-UNIID=P87306                    28.3    3.5   0.432
```

**B:** blastp **wrapper**

Figure S1: **The search results using** `blastp` **and the** `blastp` **wrapper**.

#### S1.3 The amino acid sequences that were shuffled in the null studies

Table S1: SCOP domain selected for shuffling

```
# SCOP domain selected for shuffling of length=45
>8074028 SF=3000295 SF-PDBID=2JXI_A SF-UNIID=Q88D80
MATTTLGVKLDDPTRERLKAAAQSIDRTPHWLIKQAIFNYLEKLE

# SCOP domain selected for shuffling of length=90
>8107338 SF=3000388 SF-PDBID=7PC8_A SF-UNIID=Q9NSN8
GERTVTIRRQTVGGFGLSIKGGAEHNIPVVVSKISKEQRAELSGLLFIGDAILQINGINV
RKCRHEEVVQVLRNAGEEVTLTVSFLKRAP

# SCOP domain selected for shuffling of length=175
>8107332 SF=3002300 SF-PDBID=7ECQ_A SF-UNIID=Q9D8T0
KYKCGLPQPCPEEHLSFRIVSGAANVIGPKICLEDKMLMSSVKDNVGRGLNIALVNGVSG
ELLEARAFDMWAGDVNDLLKFIRPLHEGTLVFVASYDDPATKMNEETRKLFSELGSRNAK
DLAFRDSWVFVGAKGVQNKSPFEQHMKNSKHTNKYEGWPEALEMEGCIPRRSIAG

# SCOP domain selected for shuffling of length=350
>8098538 SF=3000313 SF-PDBID=3WN6_A SF-UNIID=P17654
GQVLFQGFNWESWKENGGWYNFLMGKVDDIAAAGITHVWLPPPSHSVGEQGYMPGRLYDL
DASKYGNEAQLKSLIEAFHGKGVQVIADIVINHRTAEHKDGRGIYCLFEGGTPDSRLDWG
PHMICRDDPYGDGTGNPDTGADFAAAPDIDHLNKRVQRELIGWLDWLKMDIGFDAWRLDF
AKGYSADMAKIYIDATEPSFAVAEIWTSMANGGDGKPNYDQNAHRQELVNWVDRVGGANS
NATAFDFTTKGILNVAVEGELWRLRGEDGKAPGMIGWWPAKATTFVDNHDTGSTQHLWPF
PSDKVMQGYAYILTHPGNPCIFYDHFFDWGLKEEIERLVSIRNRQGIHPA

# SCOP domain selected for shuffling of length=700
>8076415 SF=3000970 SF-PDBID=5BV9_A SF-UNIID=A8FDC4
SNKERFLTLYHQIKSDANGYFSPEGIPYHSIETLICEAPDYGHMTTSEAYSYWLWLEVLY
GHYTRDWSKLEAAWDNMEKYIIPVNEDGNDEQPHMSAYNPSSPATYASEKPYPDQYPSQL
SGARPAGQDPIDGELKSTYGTNETYLMHWLLDVDNWYKYGNLLNPSHKAAYVNTFQRGQQ
ESVWEAIPHPSQDDKSFGKPNEGFMSLFTKENQVPAAQWRYTNATDADARAIQAIYWAKE
LGYNNSTYLDKAKKMGDFLRYGMYDKYFQTIGSGKQGNPYPGNGKGACHYLMAWYTSWGG
GLGDYANWSWRIGASHCHQGYQNPVAAYALSSDKGGLKPSSATGASDWEKTLKRQLEFYV
WLQSKEGAIAGGATNSWNGDYSAYPAGRSTFYDMAYEDAPVYHDPPSNNWFGMQAWPMER
VAELYYIFVKDGDKTSENVQMAKSCITKWVNYALDYIFIGSRPVSDEEGYFLDDQGRRIL
GGTNATVATTSAPGEFWLPGNIAWSGQPDTWNGFQSATGNPNLTAVTKDPTQDTGVLGSL
VKAFTFFAAATKLETGNYTALGVRAKDAAAQLLEVAWNYNDGVGIVTEEEREDYDRFFKK
EVYFPNGWNGTFGQGNQIPGSSTIPSDPQRGGNGVYTSFADLRPNIKQDPAWSSLESKYQ
SSFNEATGKWENGAPVFTYHRFWSQVDMATAYAEYHRLIN
```

Table S2: IID sampled sequences subject to BLAST's amino acid marginal probabilities

### we sampled the sequence for a given length from the following marginal distribution
# ['A', 'R', 'N', 'D', 'C', 'Q', 'E', 'G', 'H', 'I', 'L', 'K', 'M', 'F', 'P', 'S', 'T', 'W', 'Y', 'V']
# [0.074, 0.052, 0.045, 0.054, 0.025, 0.034, 0.054, 0.074, 0.026, 0.068, 0.099, 0.058, 0.025, 0.047, 0.039, 0.057, 0.051, 0.013, 0.032, 0.073]

### example iid sequence of length=45
>shuffled1_query_len45
QGYFDRIVNNPFTRLDQTFNSMNNCVDRYQPAAPLSMSPENSDTG

### example iid sequence of length=90
>shuffled83_query_len90
QESEVMRLAGLSTNSVEDKADKLVNFKKSSGQAERTSPATQWYFKFFKEEKSSGTIFKVA
EQIERNVALSIMLIPEIKPAFVPGTNSMVY

### example iid sequence of length=175
>shuffled1_query_len175
CKIGILDLEFNERHAADDQEWIPEKDDDLEVDGVNIATVEHTKKACEDLKRNNGLREEWL
EPETPKRLRALNATKYSSADTRKELRASNSIDDWFQERGATWVKVLAYCVMVGSRGGSAV
QTTVRRLTNIQSMKKAIAMDADALFSDSFKKDGAPEQVPGKQIIASQVLATITES

### example iid sequence of length=350
>shuffled1_query_len350
AHVLTHQGQGEVVQHKQHNKEGSQFKPALGRPLVASFELKIWPSDLLGKVLSRATLFTVY
LDRNTRNNSSGKESIVSALENRTPMSAYNVNPVVTQMATGVRPDLIRLNYGSELSGRYLQ
IRRAVSAIYNCRNICRSQKLRARHGYNKAKFIAQVTRFTFEWFQDFQQTTRCALEFIQCE
DNDSDGAFQWEDEDAEYATHLIKGRGCKWDHSPVFGIPGGLSIYQMRLWTISLIAPGLID
TNEDFRWVKFLLTARQNEGHGKIRQFPSYSTFASETDLSLNDDQSEIYTWRVDESSAVYP
VSLFCVQRSAVMQFNTFRPVLPSINNPNYMSQTFHTLYSDLSELILVHKY

### example iid sequence of length=700
>shuffled1_query_len700
ILLNVMLEGWSLAAGKKTASTPSSTVLLVQTDHDVKTHKERATLFEIGSKSASRPAVYIK
ATRWFEPADRNRTSSKAMLMVIDFKTFSKQDLDIRFRSNIMKLRFMAHTEIAWGIQWIWR
HLGQAVYFESDHSMMPPGYMLFLQTNVVELDQEPDEEGYTRLKTILANNLAQIRYENAIE
DSIIAARPRETCPSASGLSKDTLRATPFTGSDQTASENFYIKSYSDEGLDFAKLFLNTSS
LRDANAGTLFEIVALFLELELLYPKNMAVLEILPRYVKQGGAYSAGTIAGVLLDGIDETE
ASTRLYLKPLKKFIALRIGKSDIKVDVALKHESEYPFAYPVYSLINRIFTALAGQCEGDT
YANTQRYPLLGPAMGPIIVFCMAGPPVTCRAGNNALPSHETVINGKTSRLQVIDMSSSIR
QGRQPGVSRQASVLGDGIATAKMAKVVDEGDANITVRNADPLRGMNLGDKRSHSMLKGMS
LQHETGPMTLGFKMKALAAFLLAYPAYPWVNTTDRVQAIIFETKSNSQGGPNKGNRSNFF
DRAKLLSAAVAPVSGRCSKFMTMAYTMEDSGTNRLYNSLGVKVLMDMIPHSREQVYMVAM
EYTAAYLRNGQVDNYADGTGHIDQNNVEGLDEHIRSTPGRISLQFVSDRSSPGPRVRELS
ESPPYSLEHLGKTAAMDEKNQSVMEQPLRYQREGIKVVLE

#### S1.4 Supplementary Figures

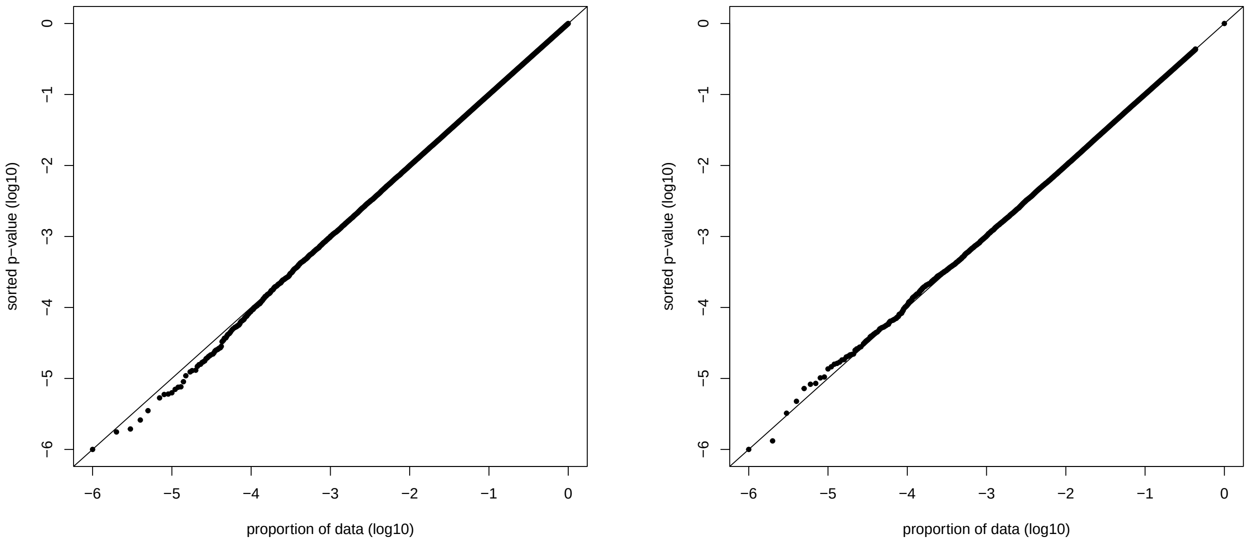


**A: Gumbel p-values applied to a Gumbel sample** **B: $SG_{50}$ p-values of a Gumbel sample**

Figure S2: **When the sample is drawn from the Gumbel distribution**. A sample of size $10^6$ was taken from the standard Gumbel distribution in order to provide a reference to our probability plots when the sample is drawn from the exact distribution rather than an approximated Gumbel. (A): The probability plot comparing the sample and the CDF of the sample's MLE fitted Gumbel. Specifically, the y-axis denotes the (log10 of the) right tail probability (p-value) of the sample points computed using the sample-fitted Gumbel. The points were sorted in decreasing order and the x-axis denotes the (log10 of the) fraction of sample points that are smaller. (B): Similar to (A) only the p-value of each sample point is the $SG_{50}$ p-value computed using the tail probability table described in Section 4.1. Specifically, each Gumbel sample value from (A) was studentized using an independent sample of size $m = 50$ from the standard Gumbel distribution, and the corresponding tail probability of the studentized value was taken from the table.

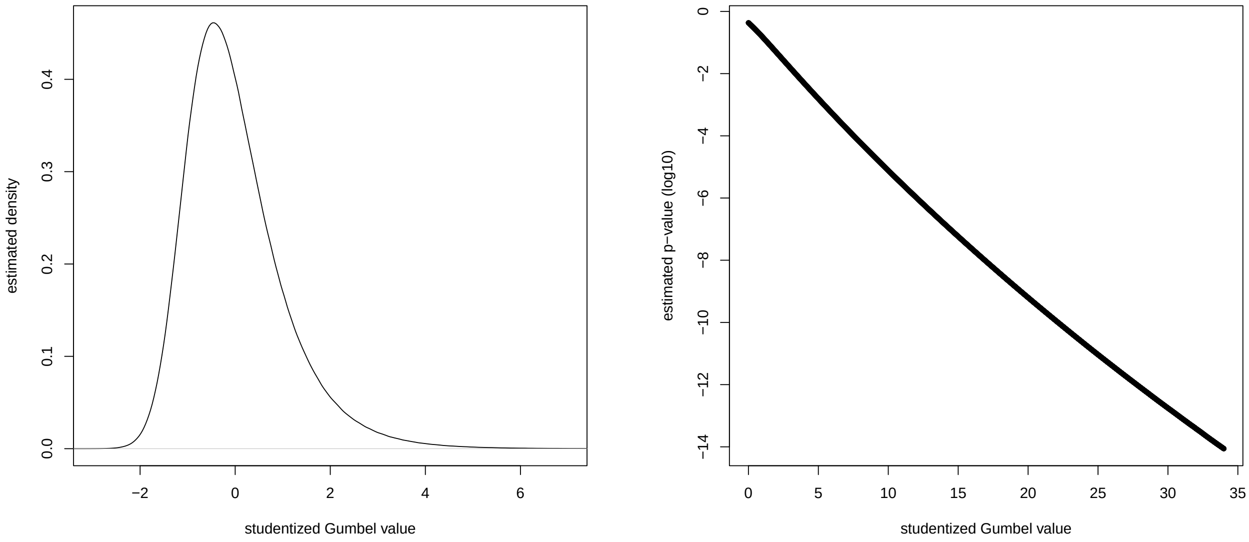


**A: Estimated $SG_{50}$ density** **B: The range of $SG_{50}$ p-values we pre-computed**

Figure S3: **The $SG_{50}$ distribution**. (A): A sample of size $10^7$ was taken from the $SG_{50}$ distribution and a density was estimated using the `R` function `density`. (B): The log10 of the right tail probabilities of the $SG_{50}$ distribution estimated via importance sampling as described in Section 4.1.

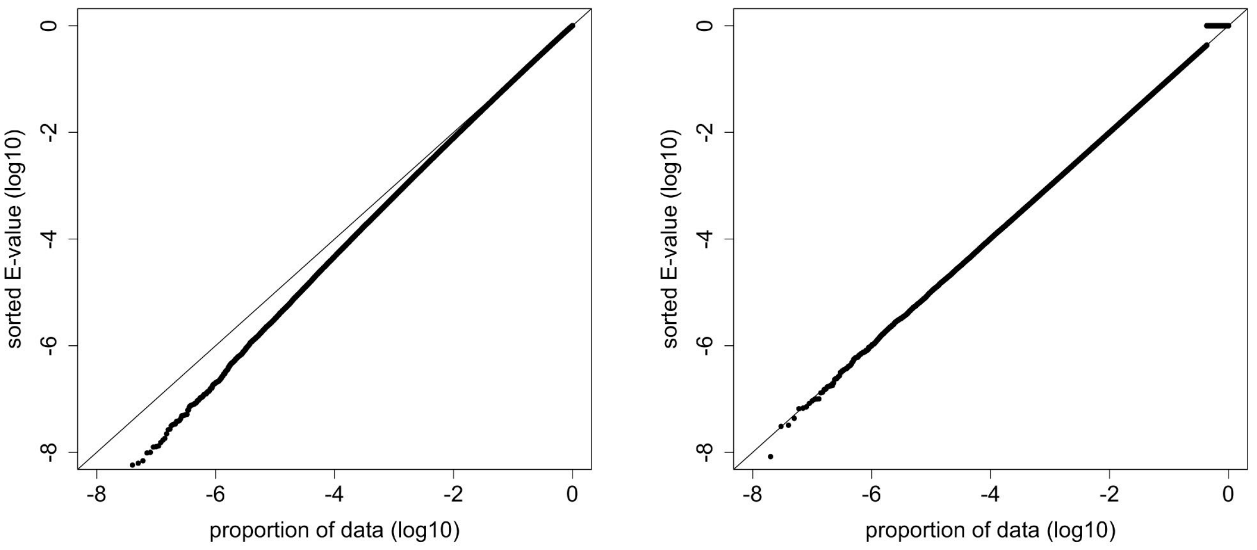


**A: MLE p-values** **B: $SG_{50}$ p-values**

Figure S4: **The MLE p-values are not valid**. (A): A sample of size $10^8$ of Gumbel MLE p-values was generated by sampling $10^8$ independent sets of $1 + m$ standard Gumbel samples: $(s_i, y_{i1}, y_{i2}, \ldots, y_{im})$ for $i = 1, 2, \ldots, 10^8$ ($m = 50$ here). Then $y_{i1}, \ldots, y_{im}$ were used to find the MLE of the Gumbel rate and location parameters, which were plugged into the Gumbel CDF to estimate the p-value of $s_i$. The probability plot above shows these MLE p-values are invalid. Specifically, for example, a p-value of $10^{-5}$ or smaller was observed at a rate of $2.6 \cdot 10{-}5$, i.e., 2.6 times more often than expected (binomial test p-value of 0). (B): The same $10^8$ sampled sets that were used in (A) were used here to compute the $SG_{50}$ p-value of each $s_i$. Specifically, instead of estimating the Gumbel parameters from the $y_{i\bullet}$ sample, it was used to studentize $s_i$ and the p-value was taken from our tabulated right tail of $SG_{50}$. The $SG_{50}$ p-values of the same sample used in (A) appear to be valid.

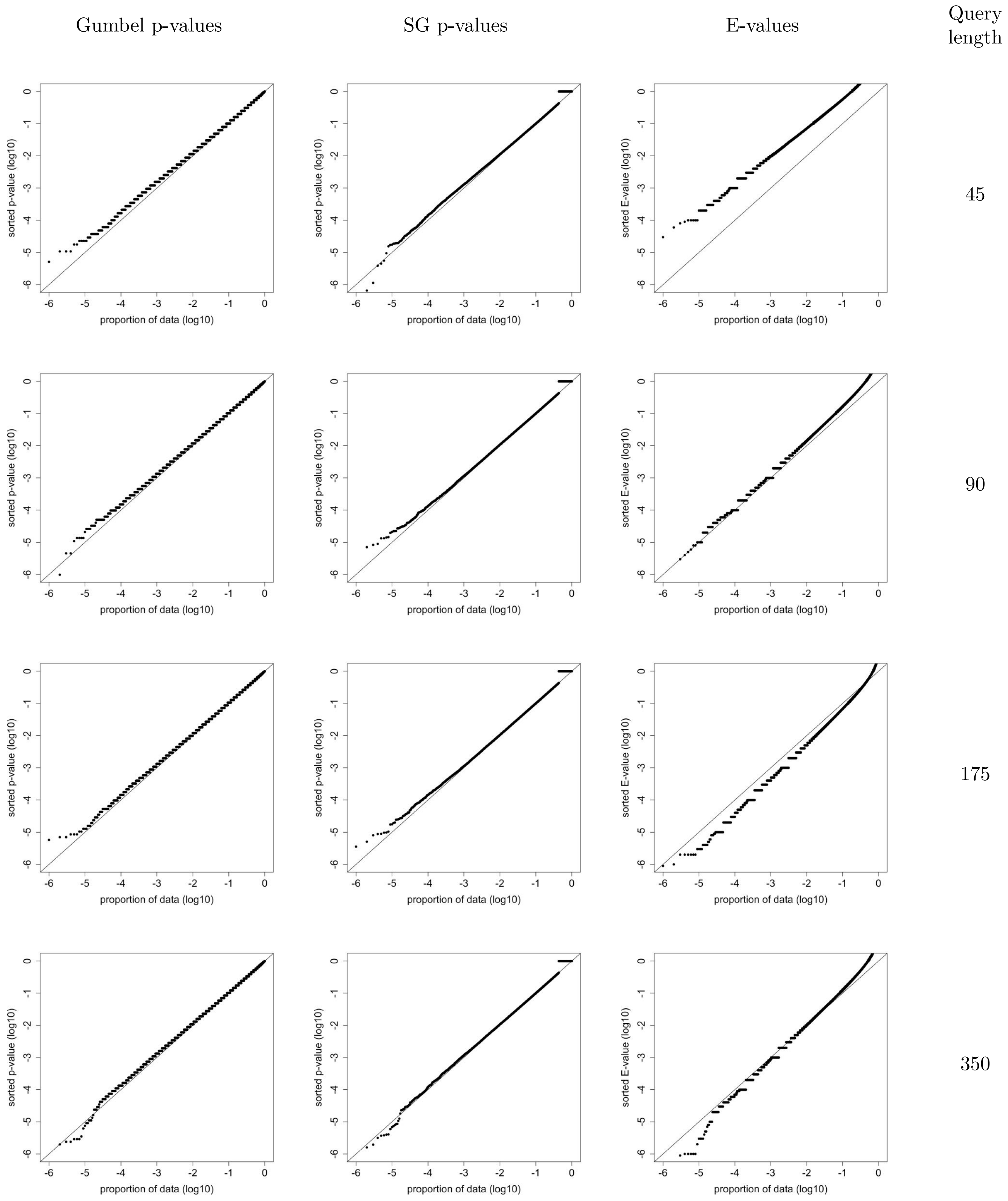


Figure S5: **Using BLAST to search $10^6$ shuffled queries against the Swiss-Prot database with BLOSUM45 (14,2)**. Five domains from the SCOP database of lengths 45, 90, 175, and 350 were selected, and $10^6$ shuffles of each were blasted against Swiss-Prot using BLOSUM45 (14,2) and the database-wide optimal local alignment (raw) score and E-values were noted. Each of the probability plots specifies the (log10 of the) fraction of the shuffles (x-axis) for which the optimal alignment p-value/E-value is better than the (log10 of the) reported optimal p-value/E-value (y-axis). Note that the y-axis was constrained to match the range of the x-axis (log10 of the number of shuffles). BLAST's E-values here are at times overly conservative, e.g., only 0.1% of the length 45 E-values are $\leq 0.01$, and at others too liberal, e.g., 7.7% of the length 175 E-values are $\leq 0.05$ (binomial test p-value of 0). In contrast, the $SG_{50}$ p-values are valid and better calibrated for all query lengths.

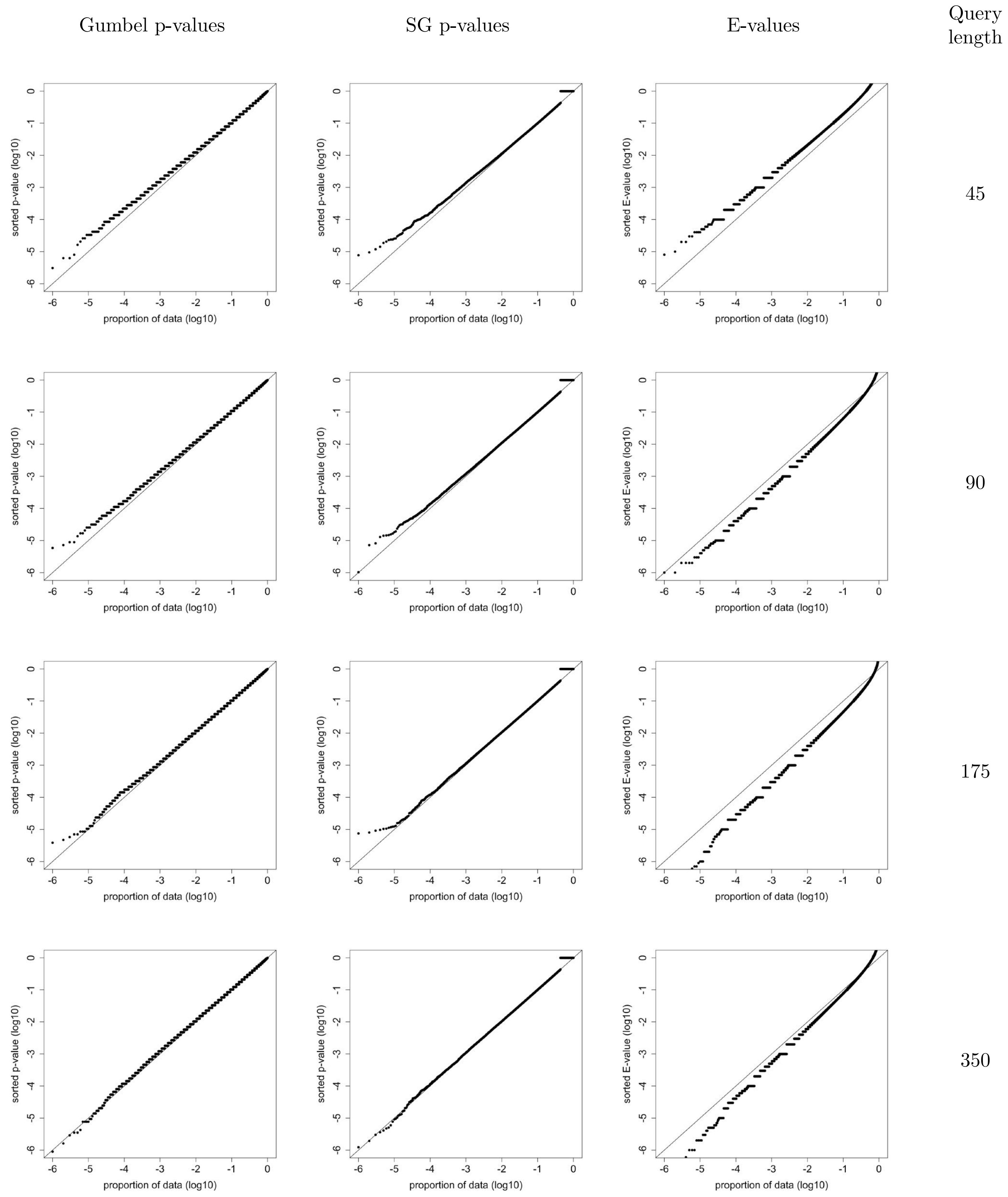


Figure S6: **Using BLAST to search $10^6$ shuffled queries against the SCOP database with BLOSUM45 (14,2)**. Same as Figure S5 only searching the SCOP database (rather than Swiss-Prot). Note that the y-axis was constrained to match the range of the x-axis (log10 of the number of shuffles). While not as conservative as when searching the length 45 queries Swiss-Prot, searching SCOP with the same BLOSUM45 (14,2) is visibly more liberally biased, e.g., for length 175 11% of the length 175 E-values are $\leq 0.05$ (p-value of 0). In contrast, the $SG_{50}$ p-values are valid and better calibrated for all query lengths.

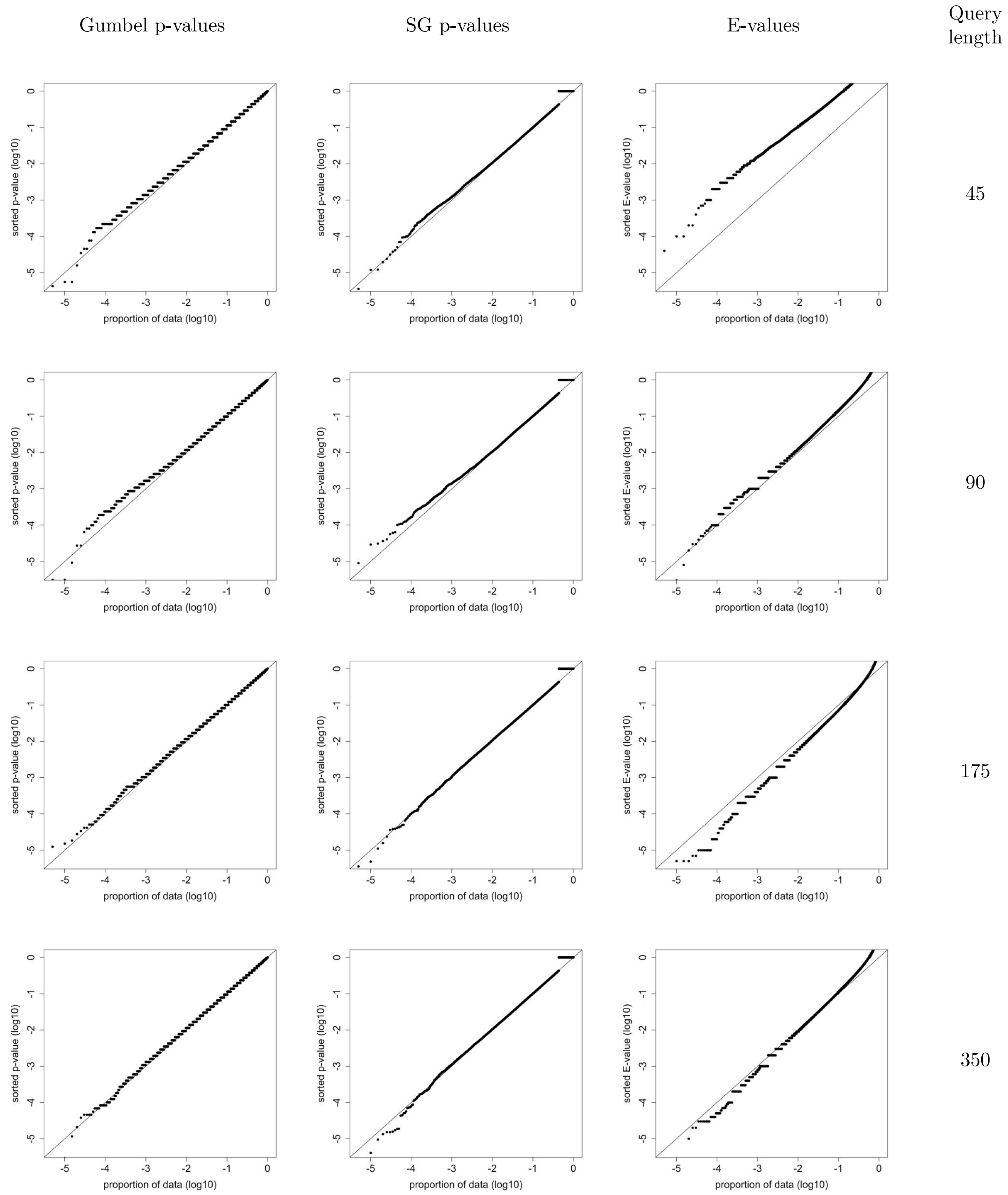


Figure S7: **Using BLAST to search 200,000 shuffled *iid-sampled* queries against the Swiss-Prot database using BLOSUM45 (14,2)**. Similar to Figure S5 but blasting (against Swiss-Prot using BLOSUM45 (14,2)) 200,000 random shuffles of five queries of length 45, 90, 175 and 350 residues, which were randomly drawn according to an iid process. Note that the y-axis was constrained to match the range of the x-axis (log10 of the number of shuffles). Qualitatively the results are the same as when searching the same Swiss-Prot using the $10^6$ shuffles of the four selected SCOP domains (Figure S5).

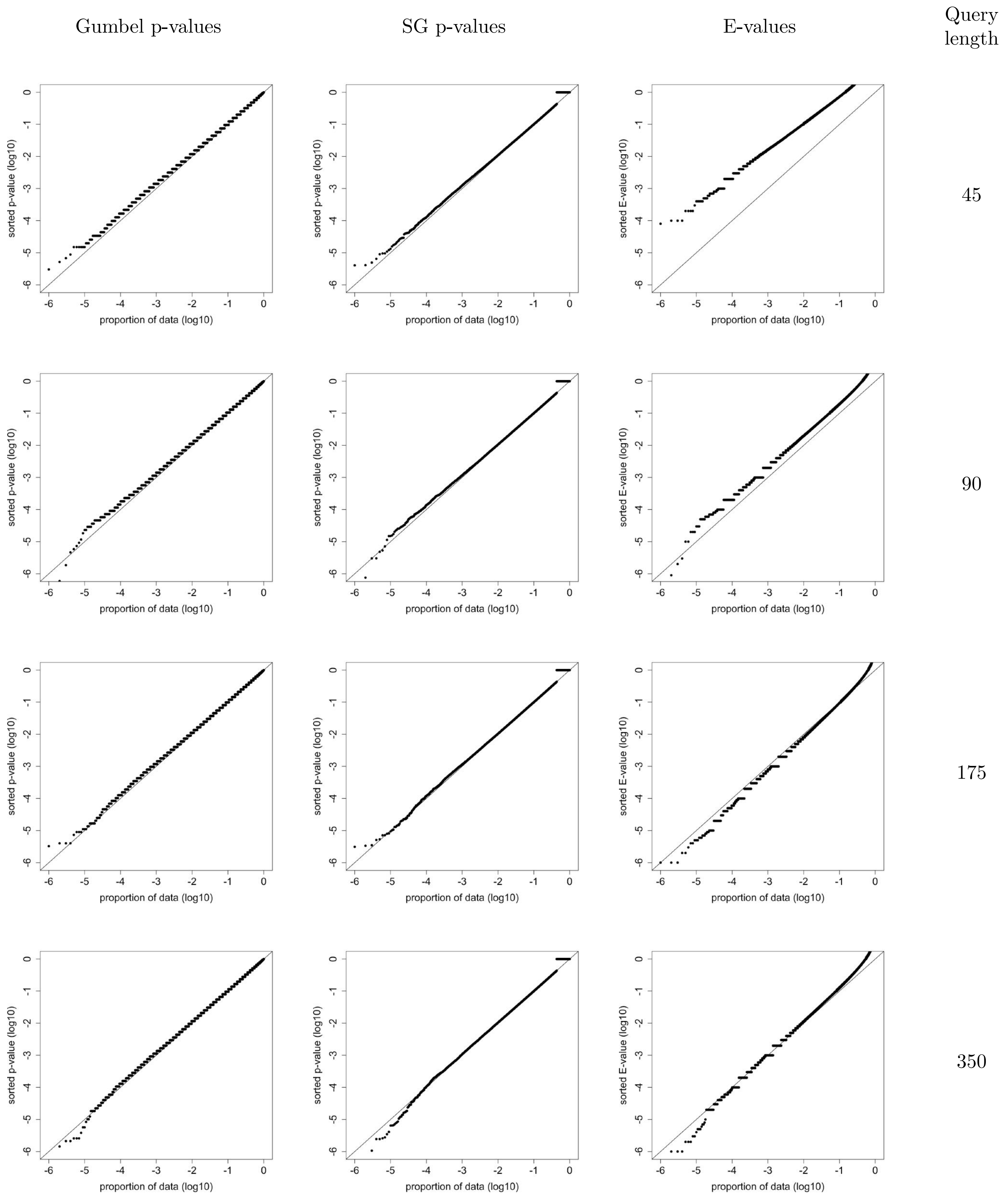


Figure S8: **Using BLAST to search $10^6$ shuffled queries against the Swiss-Prot database with BLOSUM50 (13,2)**. Same as Figure S5 only searching using BLOSUM50 (13,2) (instead of BLOSUM45). The E-values here are at times overly conservatives, e.g., only 0.06% of the length 45 E-values are $\leq 0.01$, and at others too liberal, e.g., 5.7% of the length 175 E-values are $\leq 0.05$ (binomial test p-value of $5 \cdot 10^{-197}$). In contrast, the $SG_{50}$ p-values are valid and better calibrated for all query lengths.

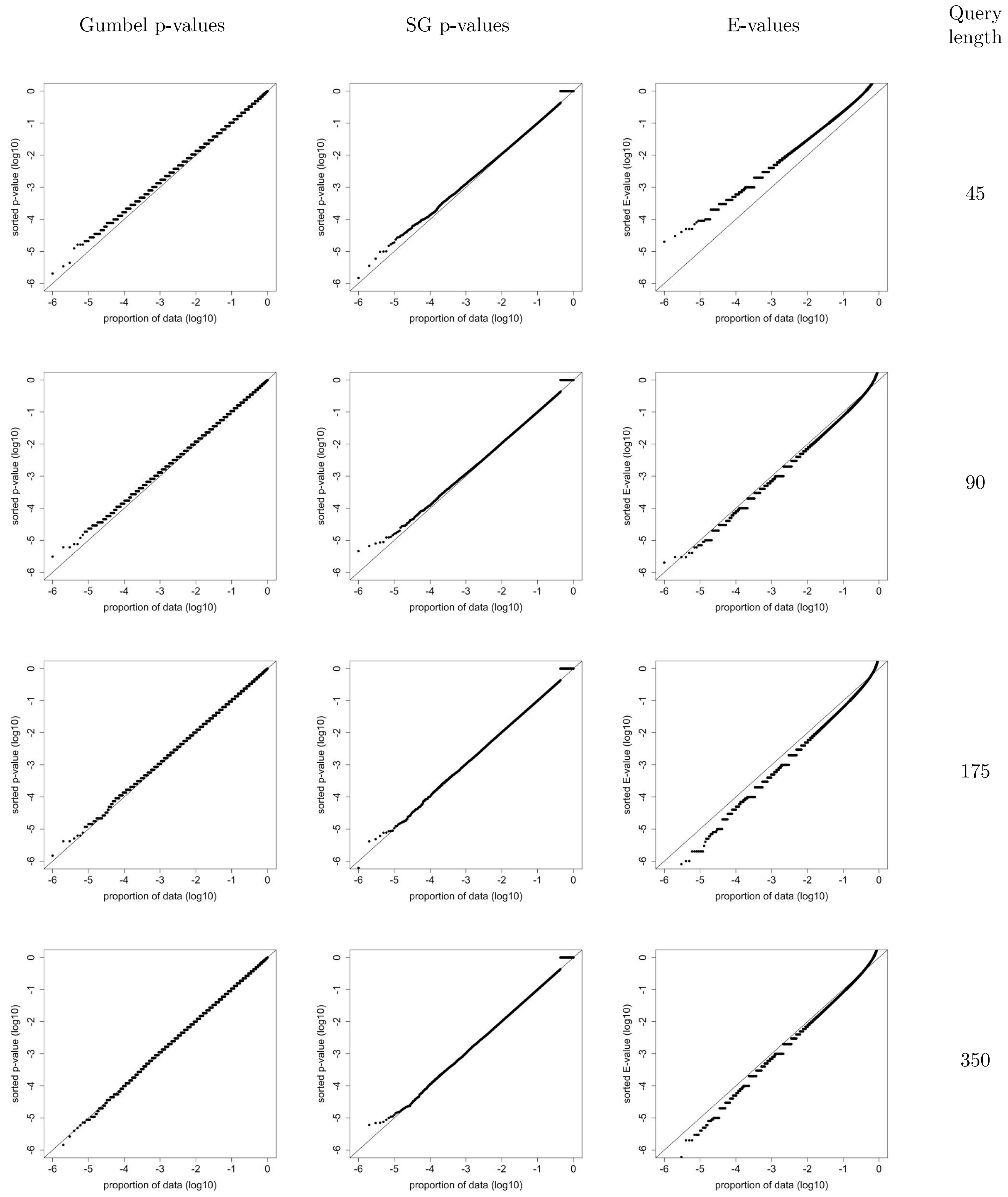


Figure S9: **Using BLAST to search $10^6$ shuffled queries against the SCOP database with BLOSUM50 (13,2).** Same as Figure S6 except searching using BLOSUM50 (13,2). For the length 175 queries there are more than 3 times as many E-values $\leq 0.001$ as expected (p-value of 0).

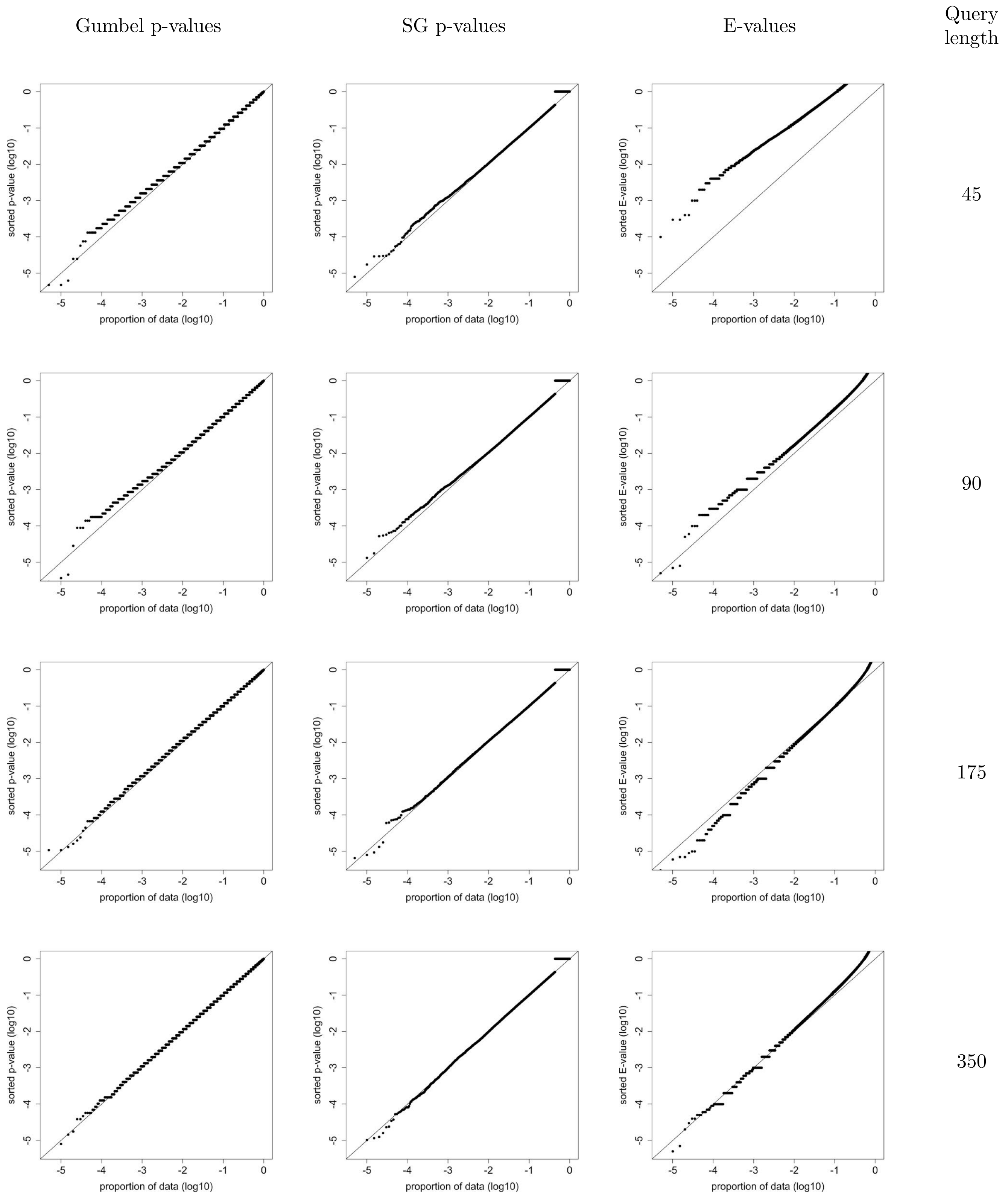


Figure S10: **Using BLAST to search 200,000 shuffled *iid-sampled* queries against the Swiss-Prot database using BLOSUM50 (13,2)**. Same as Figure S7 only using BLOSUM50 (13,2).

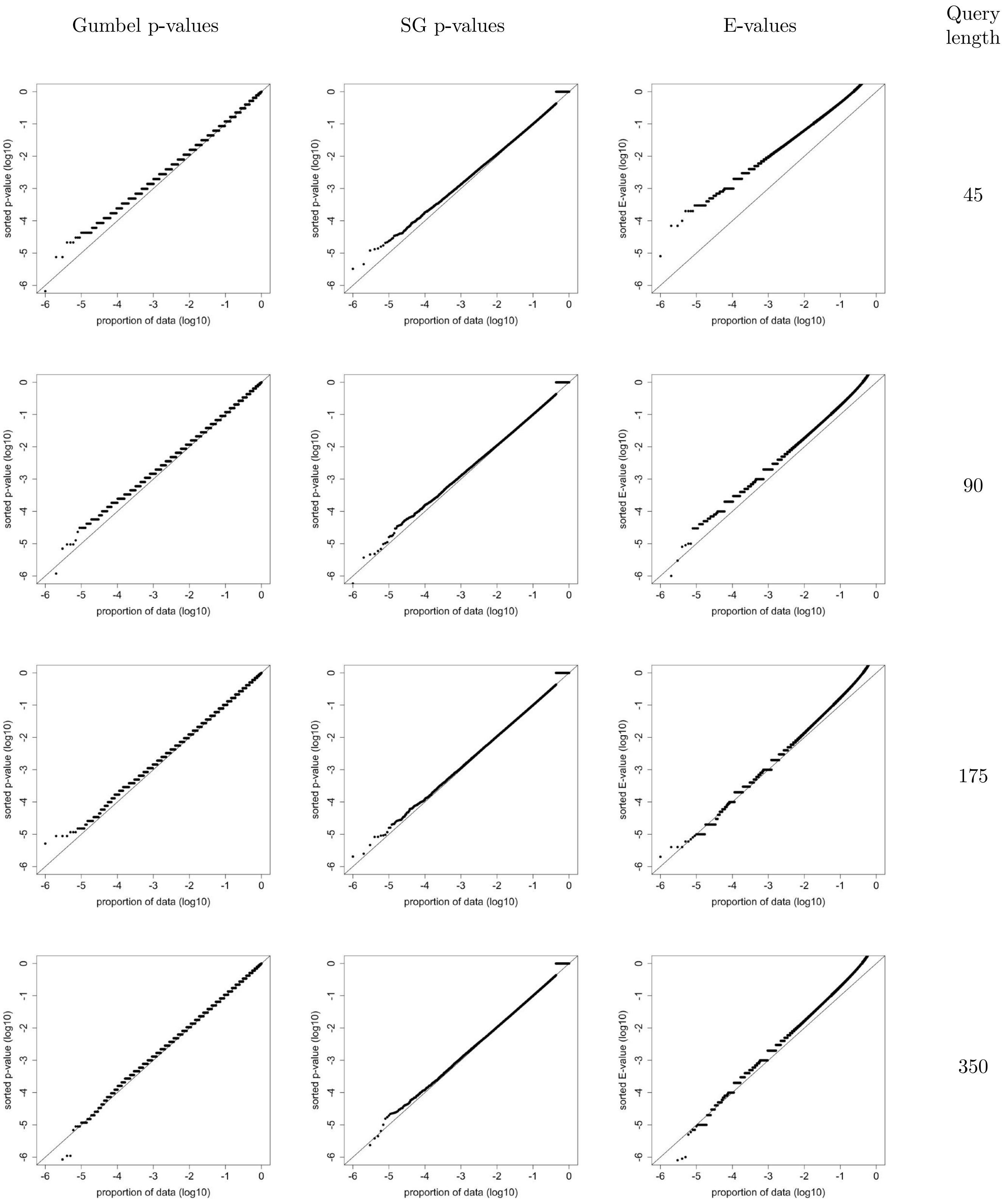


Figure S11: **Using BLAST to search $10^6$ shuffled queries against the Swiss-Prot database with BLOSUM62 (11,1)**. Same as Figure S5 except searching using BLOSUM62 (11,1). Only 4 of the length 45 shuffles and 6 of the length 90 shuffles have an E-value $\leq 10^{-4}$ instead of the expected 100 in each case. At the same time there 22% more length 175 E-values that are $\leq 0.001$ than expected (p-value of $1.7 \cdot 10^{-11}$).

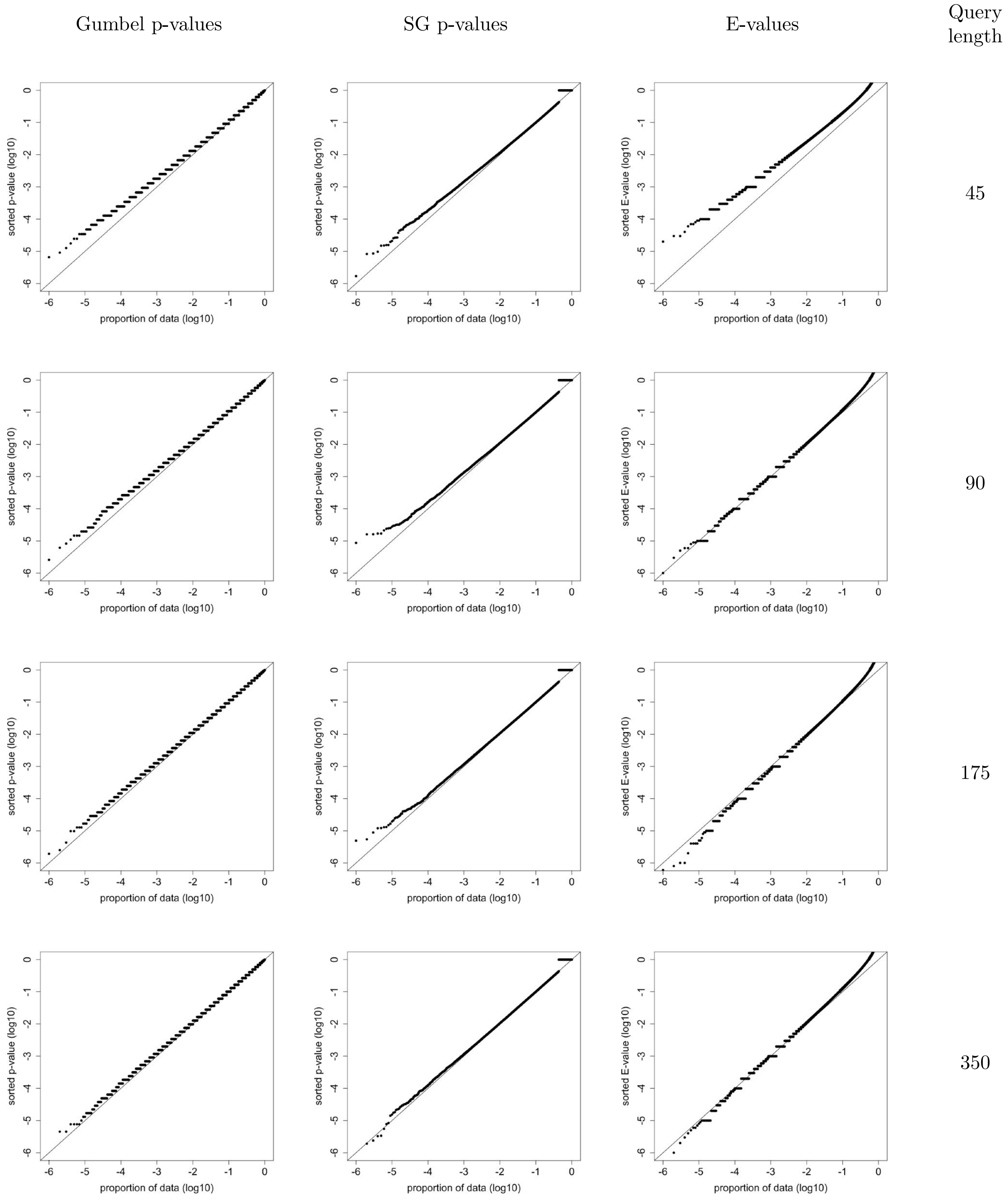


Figure S12: **Using BLAST to search $10^6$ shuffled queries against the SCOP database with BLOSUM62 (11,1).** Same as Figure S6 except searching using BLOSUM62 (11,1). For the length 175 shuffles there are 78% more E-values $\leq 0.001$ than expected (p-value of $5 \cdot 10^{-109}$).

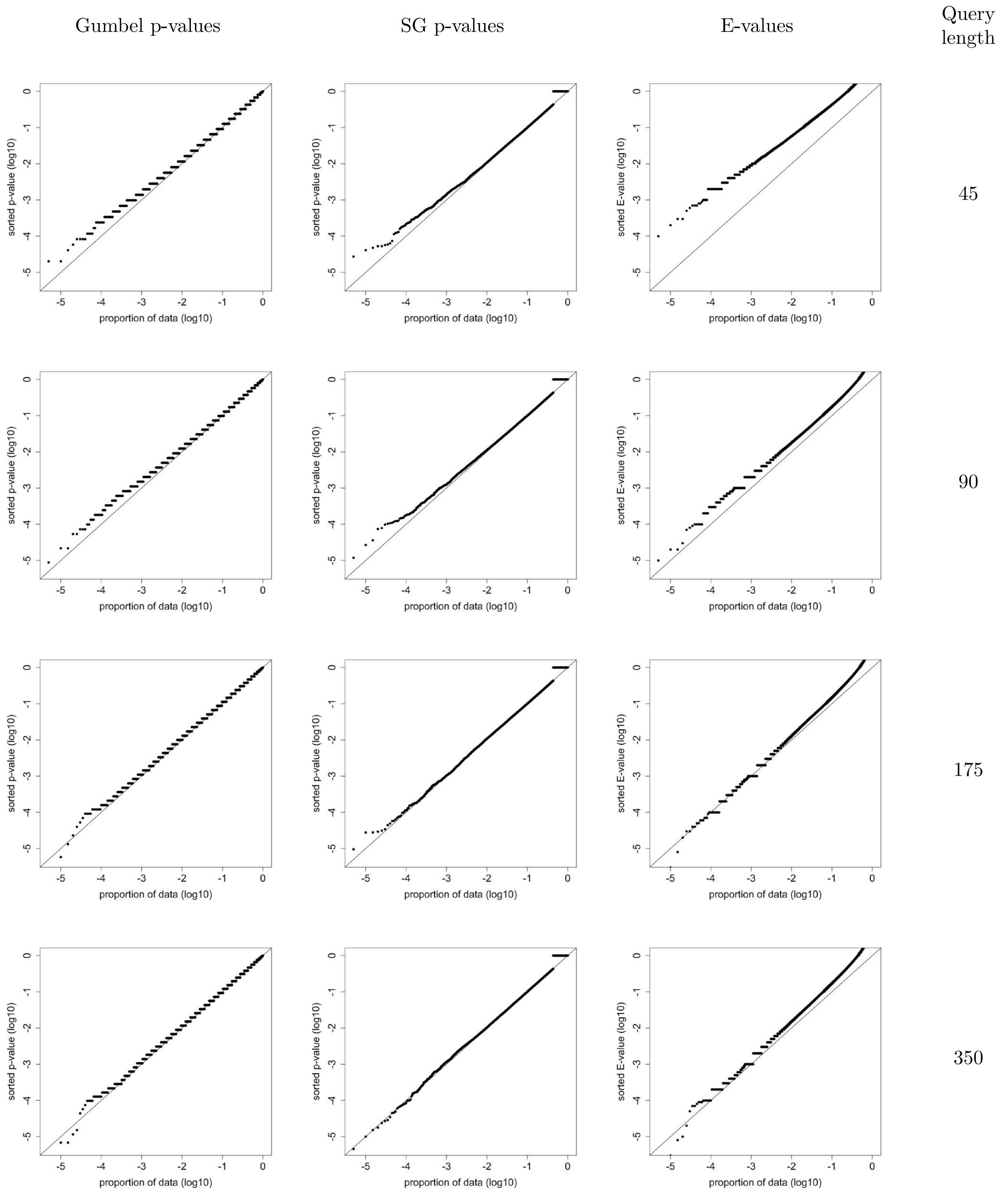


Figure S13: **Using BLAST to search 200,000 shuffled *iid-sampled* queries against the Swiss-Prot database using BLOSUM62 (11,1)**. Same as Figure S7 only using BLOSUM62 (11,1).

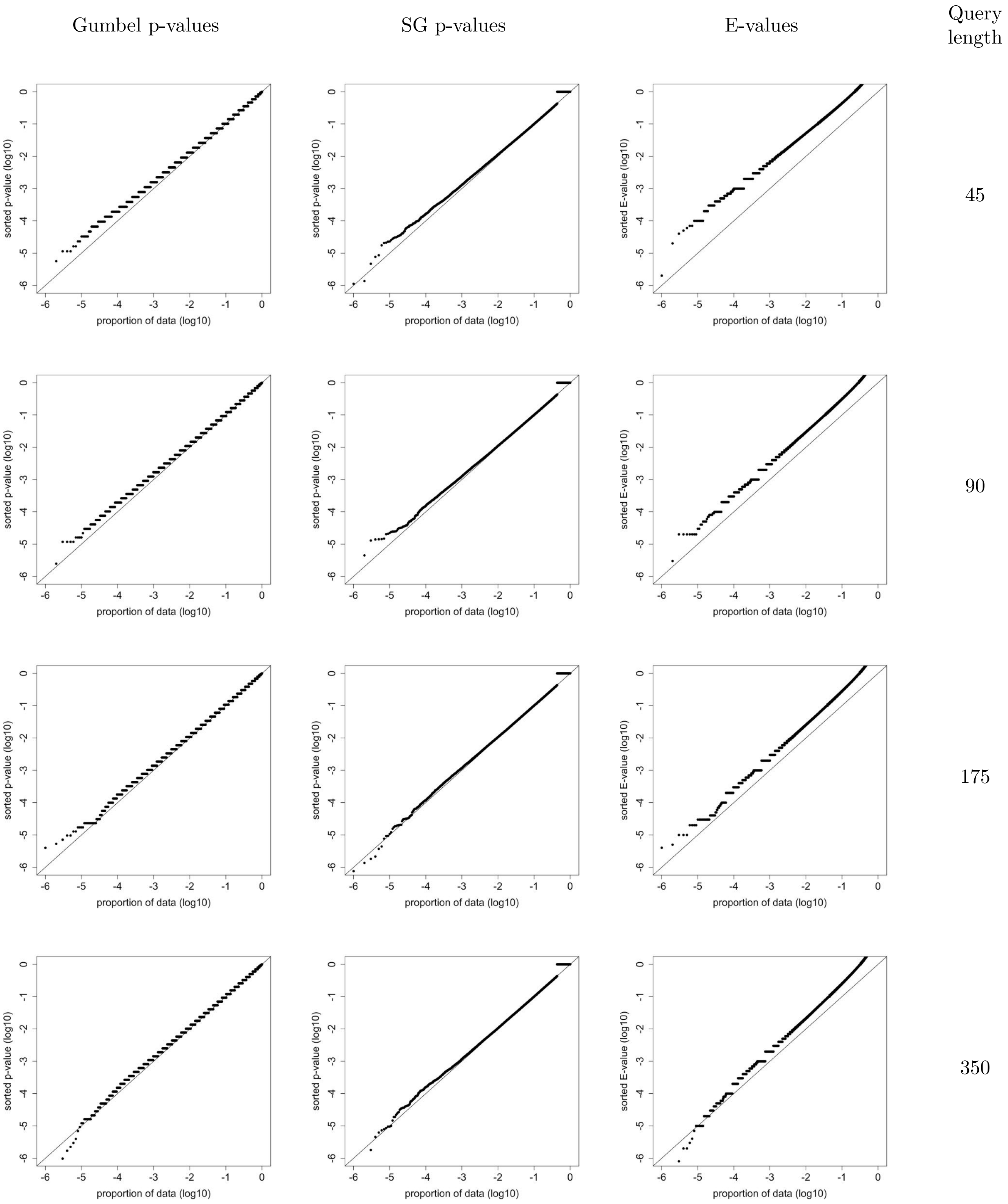


Figure S14: **Using BLAST to search $10^6$ shuffled queries against the Swiss-Prot database with BLOSUM80 (10,1)**. Same as Figure S5 except searching using BLOSUM80.

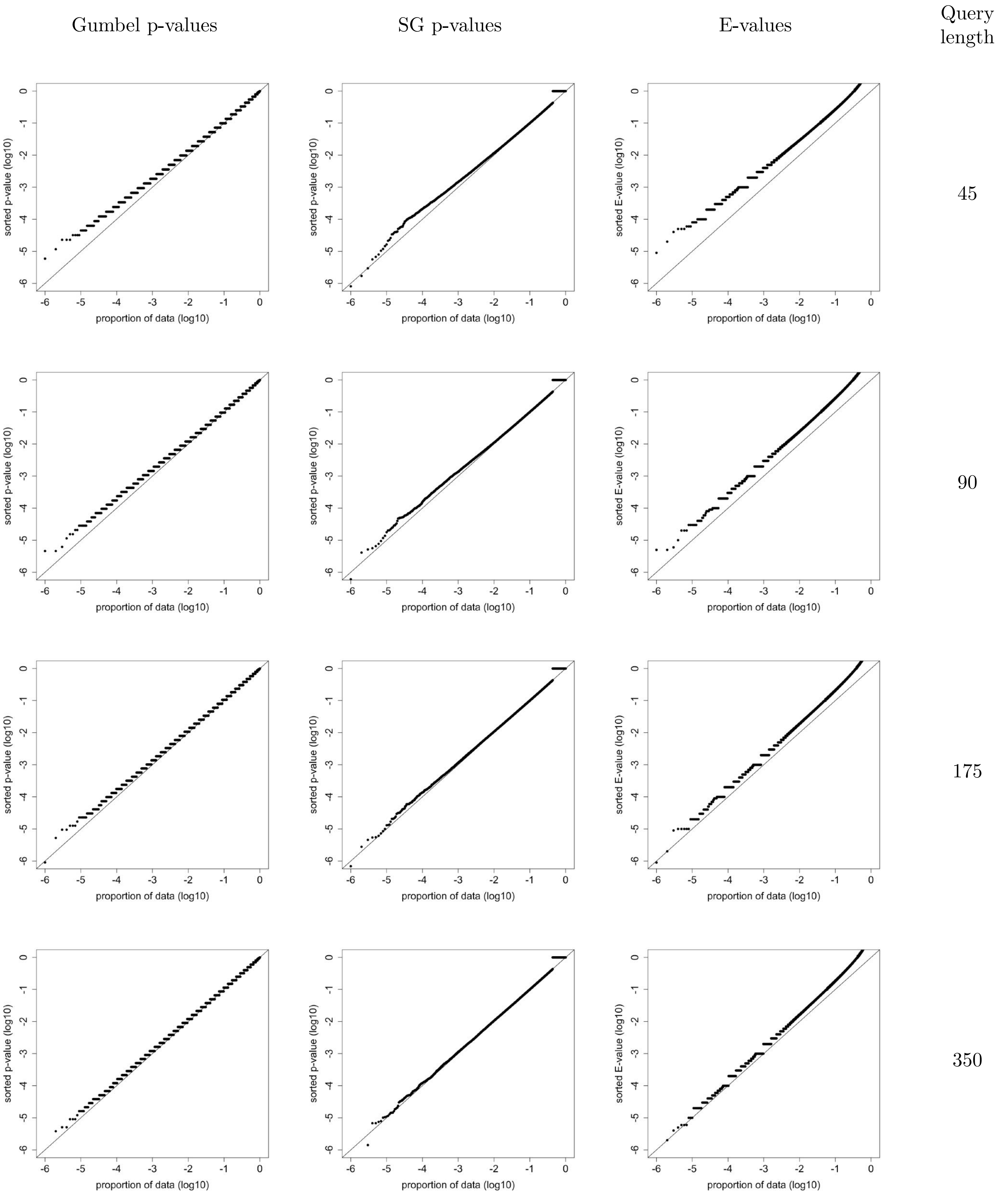


Figure S15: **Using BLAST to search $10^6$ shuffled queries against the SCOP database with BLOSUM80 (10,1)**. Same as Figure S6 except searching using BLOSUM80 (10,1).

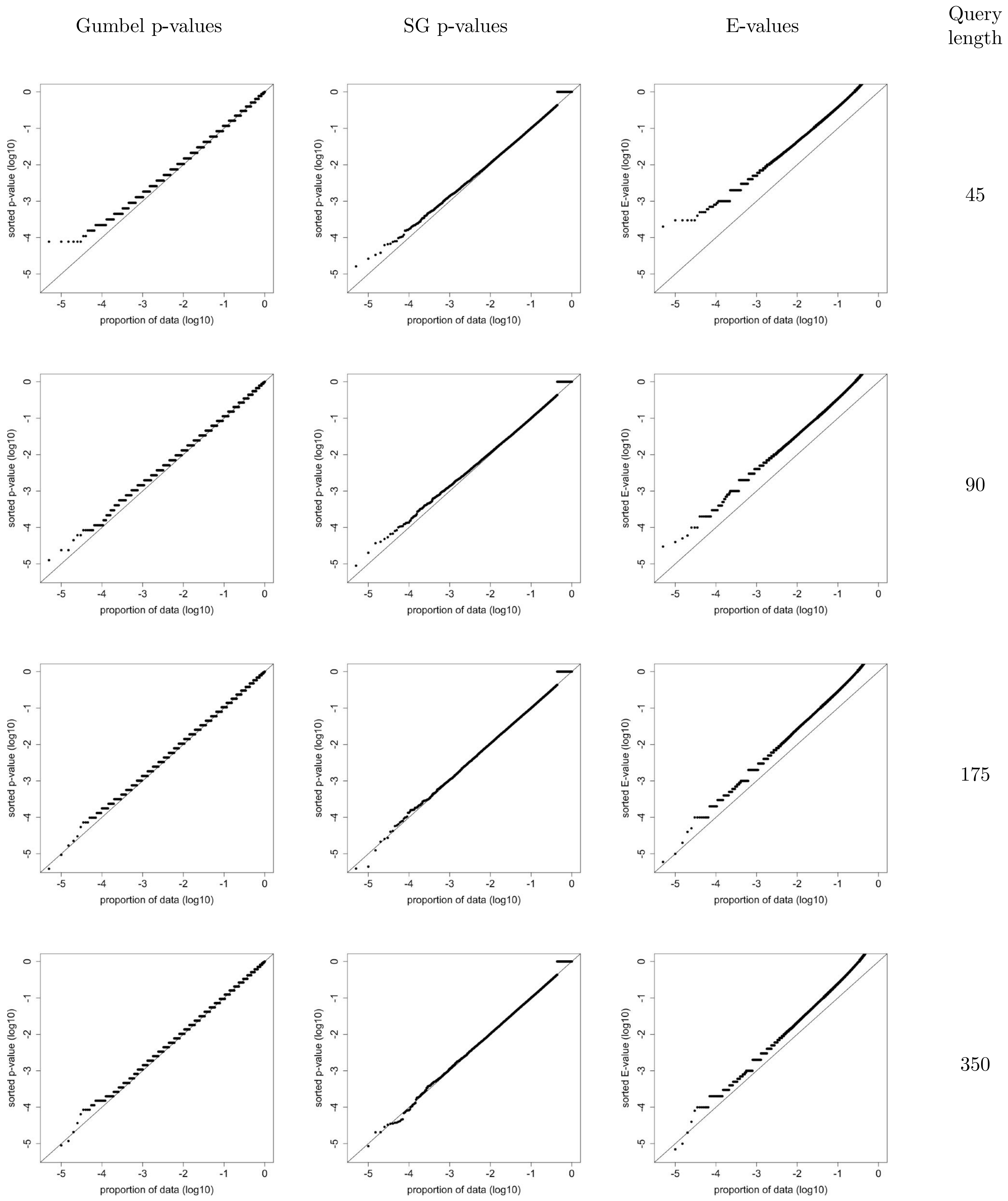


Figure S16: **Using BLAST to search 200,000 shuffled *iid-sampled* queries against the Swiss-Prot database using BLOSUM80 (10,1)**. Same as Figure S7 except using BLOSUM80 (10,1).

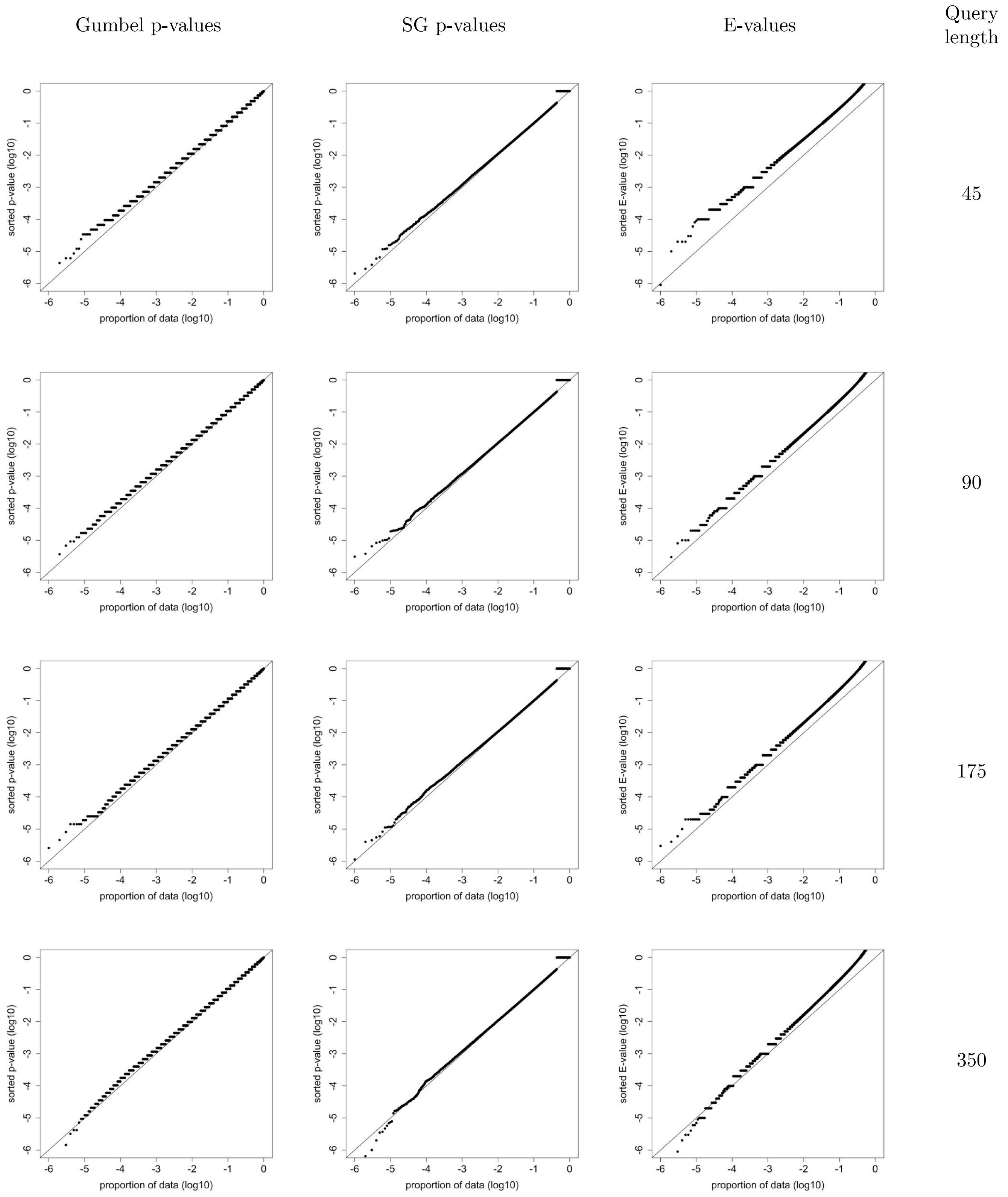


Figure S17: **Using BLAST to search $10^6$ shuffled queries against the Swiss-Prot database with BLOSUM90 (10,1)**. Same as Figure S5 except searching using BLOSUM90.

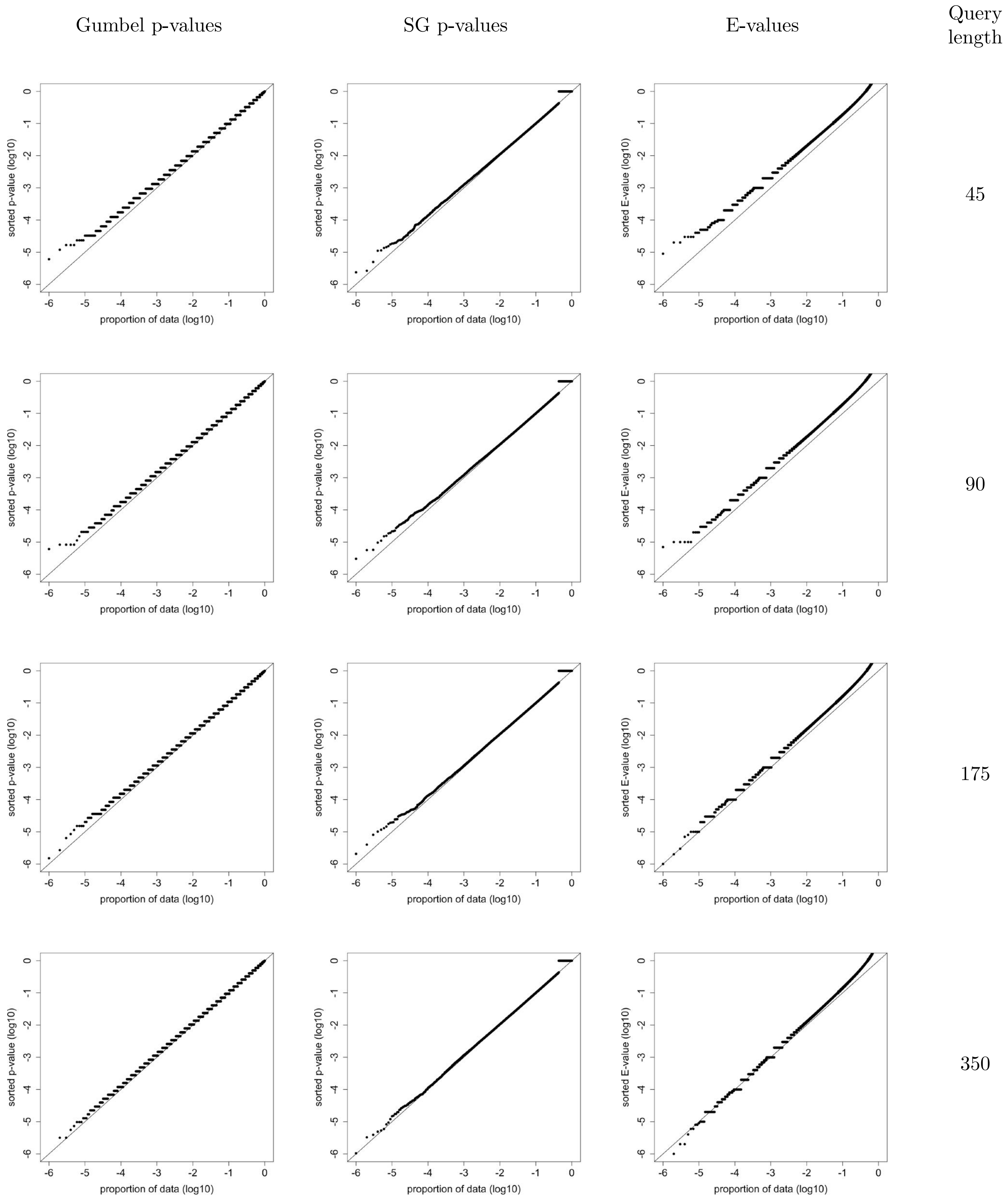


Figure S18: **Using BLAST to search $10^6$ shuffled queries against the SCOP database with BLOSUM90 (10,1)**. Same as Figure S6 except searching using BLOSUM90 (10,1).

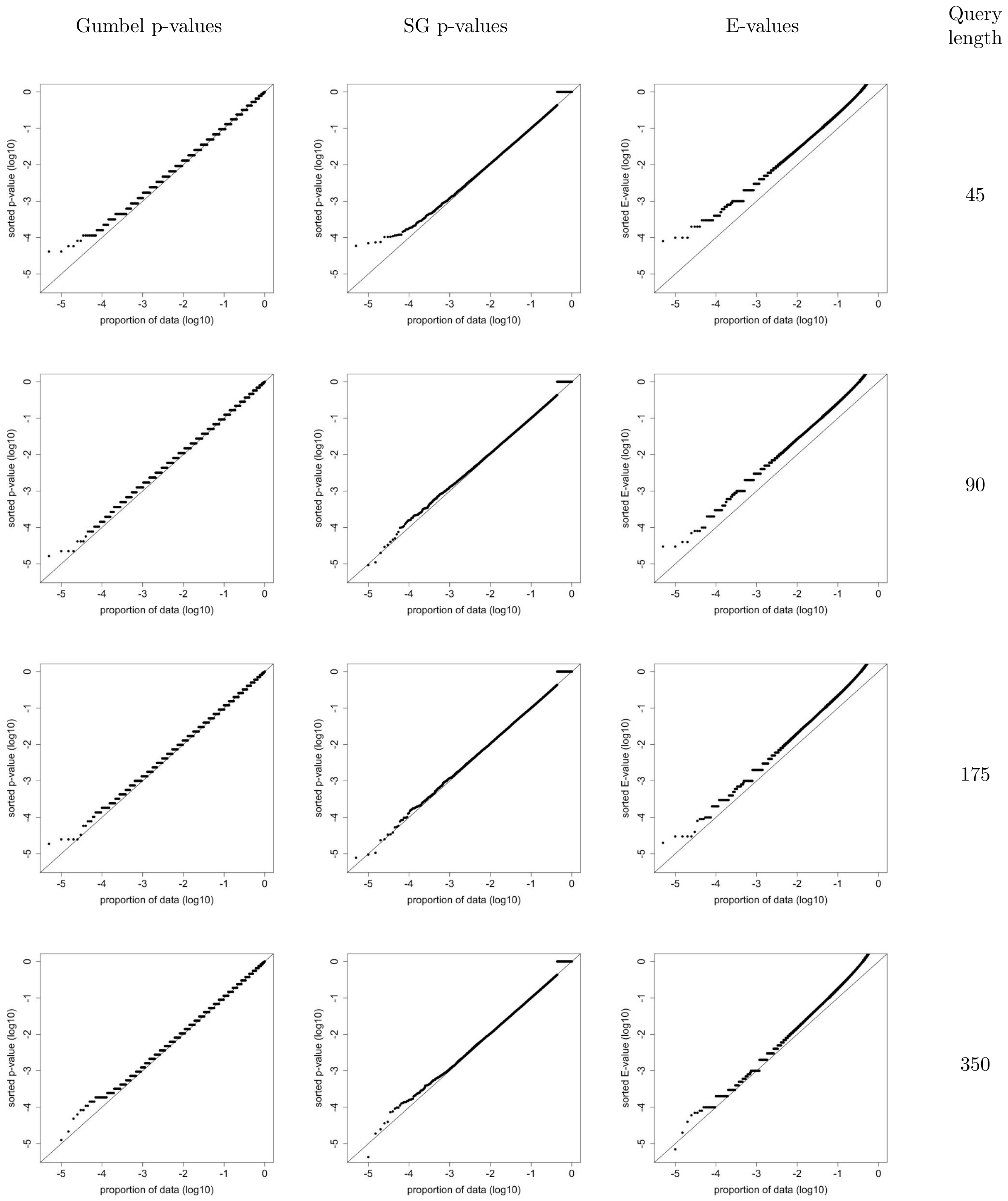


Figure S19: **Using BLAST to search 200,000 shuffled *iid-sampled* queries against the Swiss-Prot database using BLOSUM90 (10,1)**. Same as Figure S7 except using BLOSUM90 (10,1).

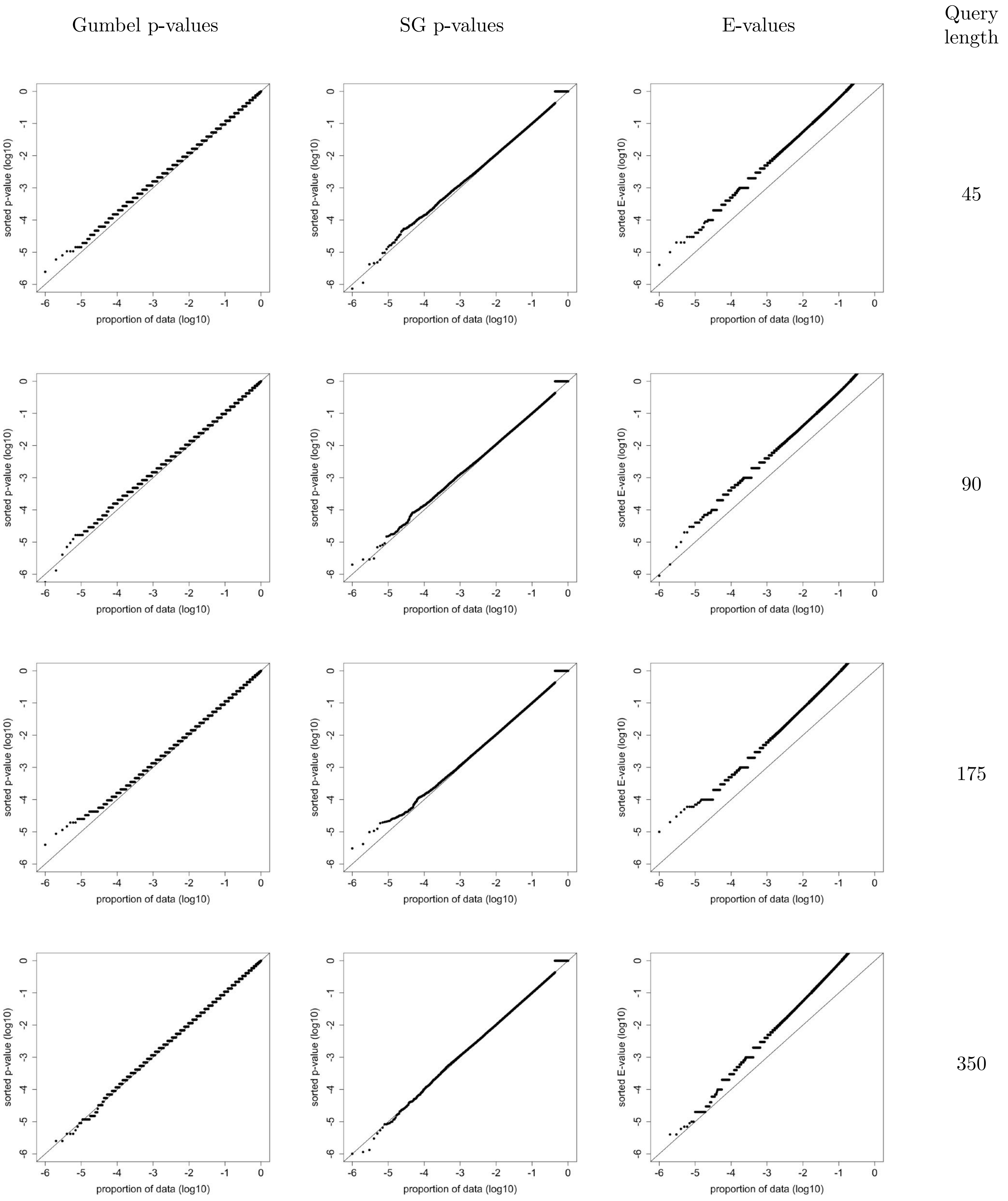


Figure S20: **Using BLAST to search $10^6$ shuffled queries against the Swiss-Prot database with PAM30 (9,1).** Same as Figure S5 except searching using PAM30 (9,1).

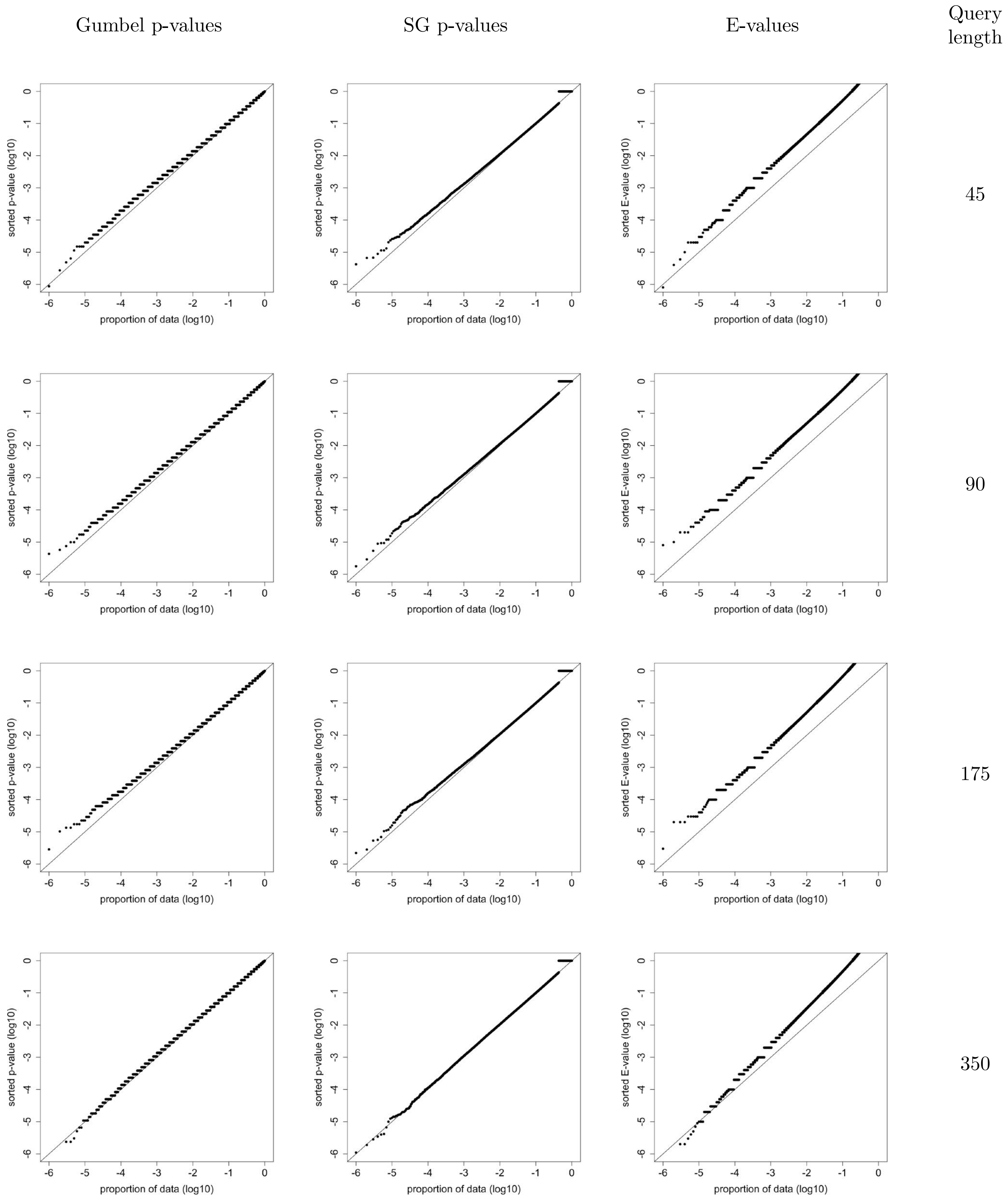


Figure S21: **Using BLAST to search** $10^6$ **shuffled queries against the SCOP database with PAM30 (9,1)**. Same as Figure S6 except searching using PAM30 (9,1).

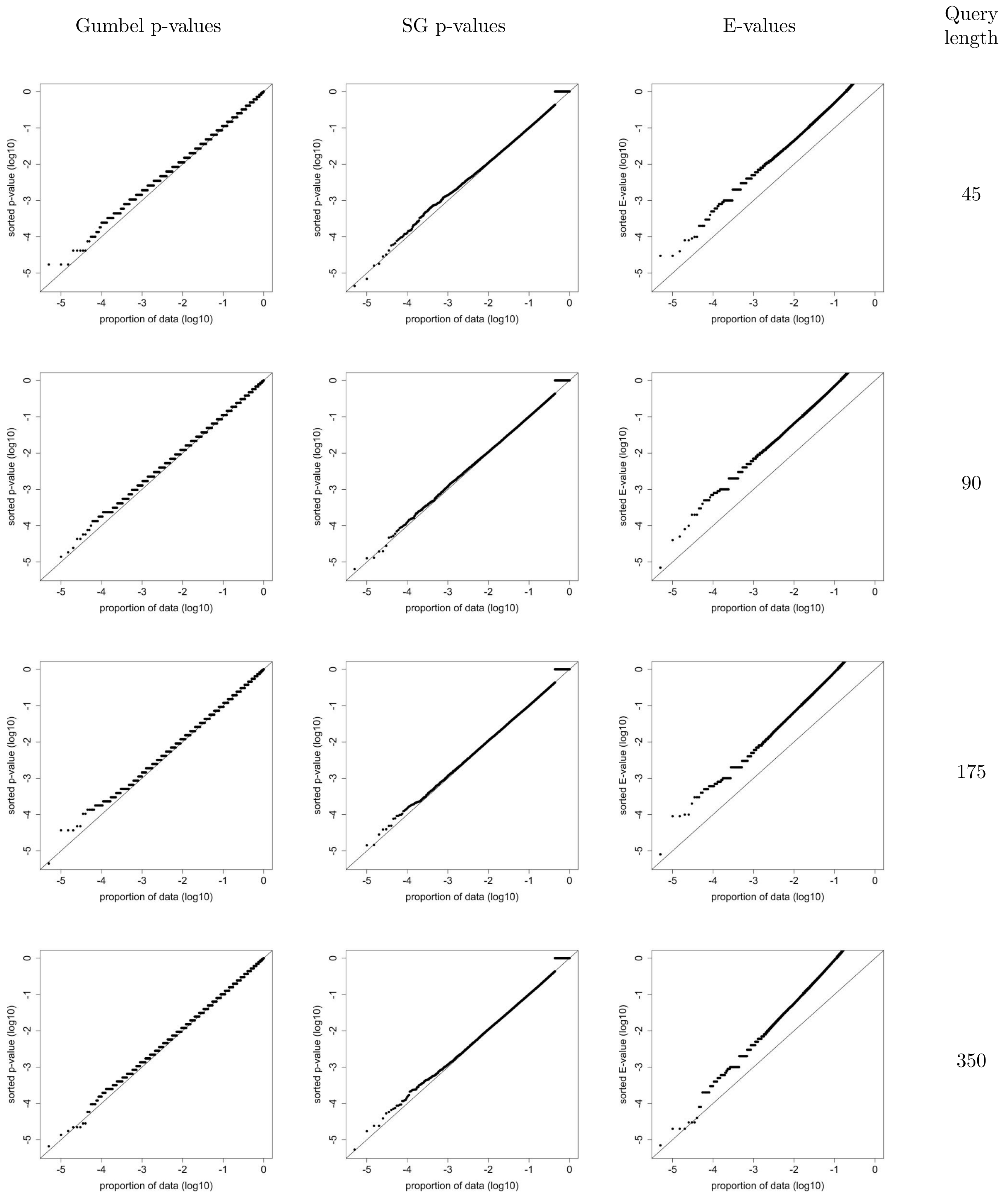


Figure S22: **Using BLAST to search 200,000 shuffled *iid-sampled* queries against the Swiss-Prot database using PAM30 (9,1)**. Same as Figure S7 except using PAM30 (9,1).

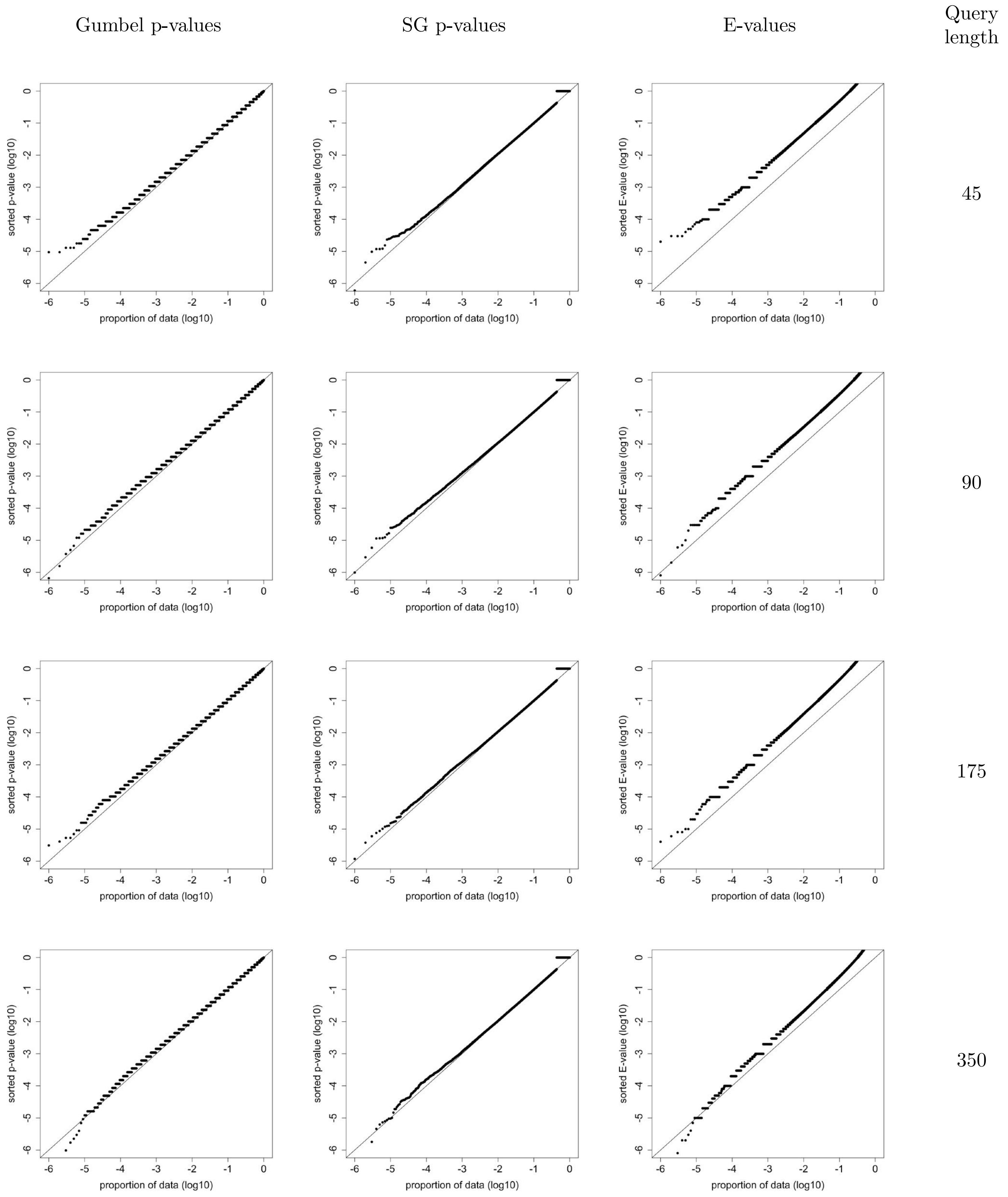


Figure S23: **Using BLAST to search $10^6$ shuffled queries against the Swiss-Prot database with PAM70 (10,1).** Same as Figure S5 except searching using PAM70.

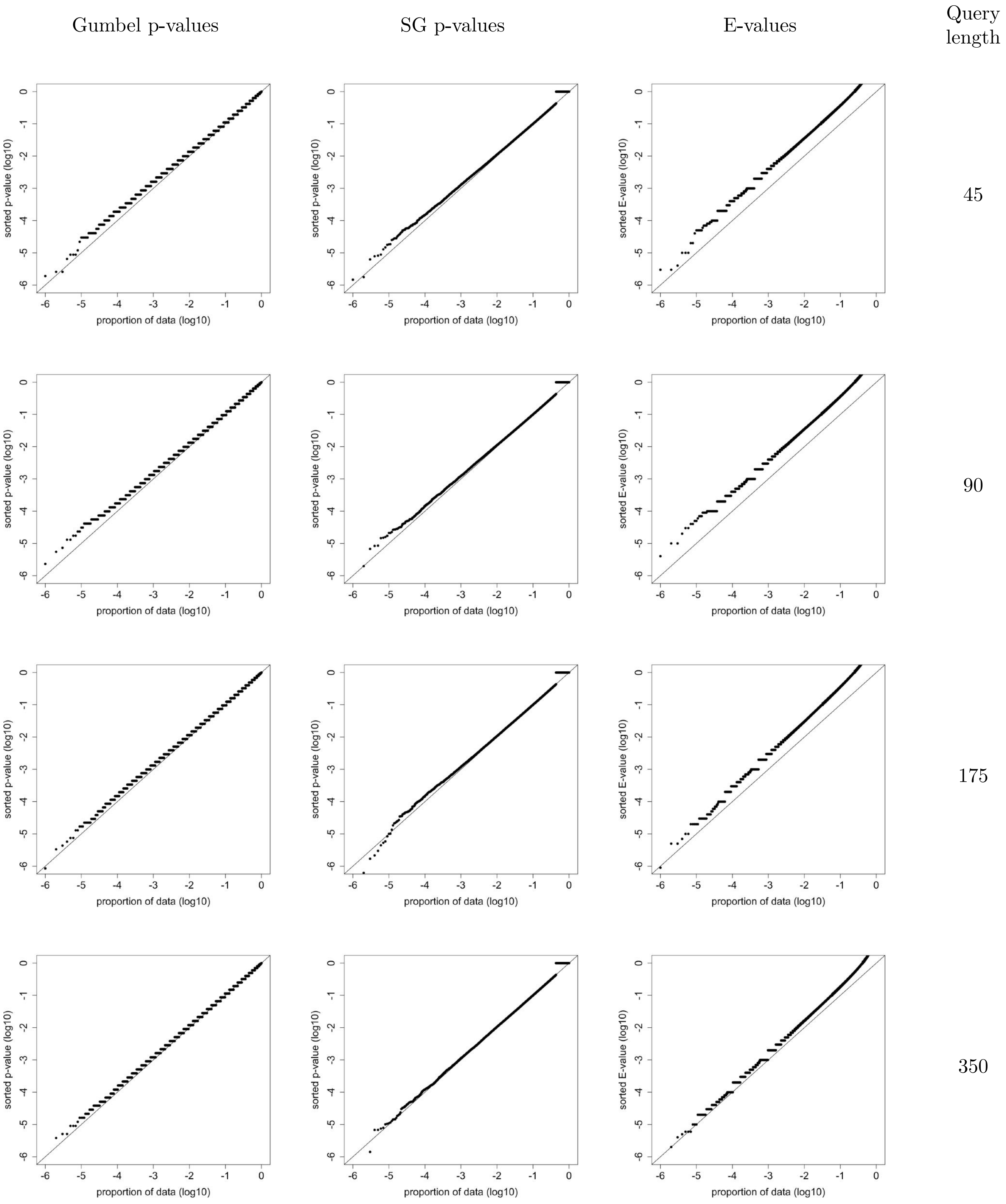


Figure S24: **Using BLAST to search $10^6$ shuffled queries against the SCOP database with PAM70 (10,1)**. Same as Figure S6 except searching using PAM70.

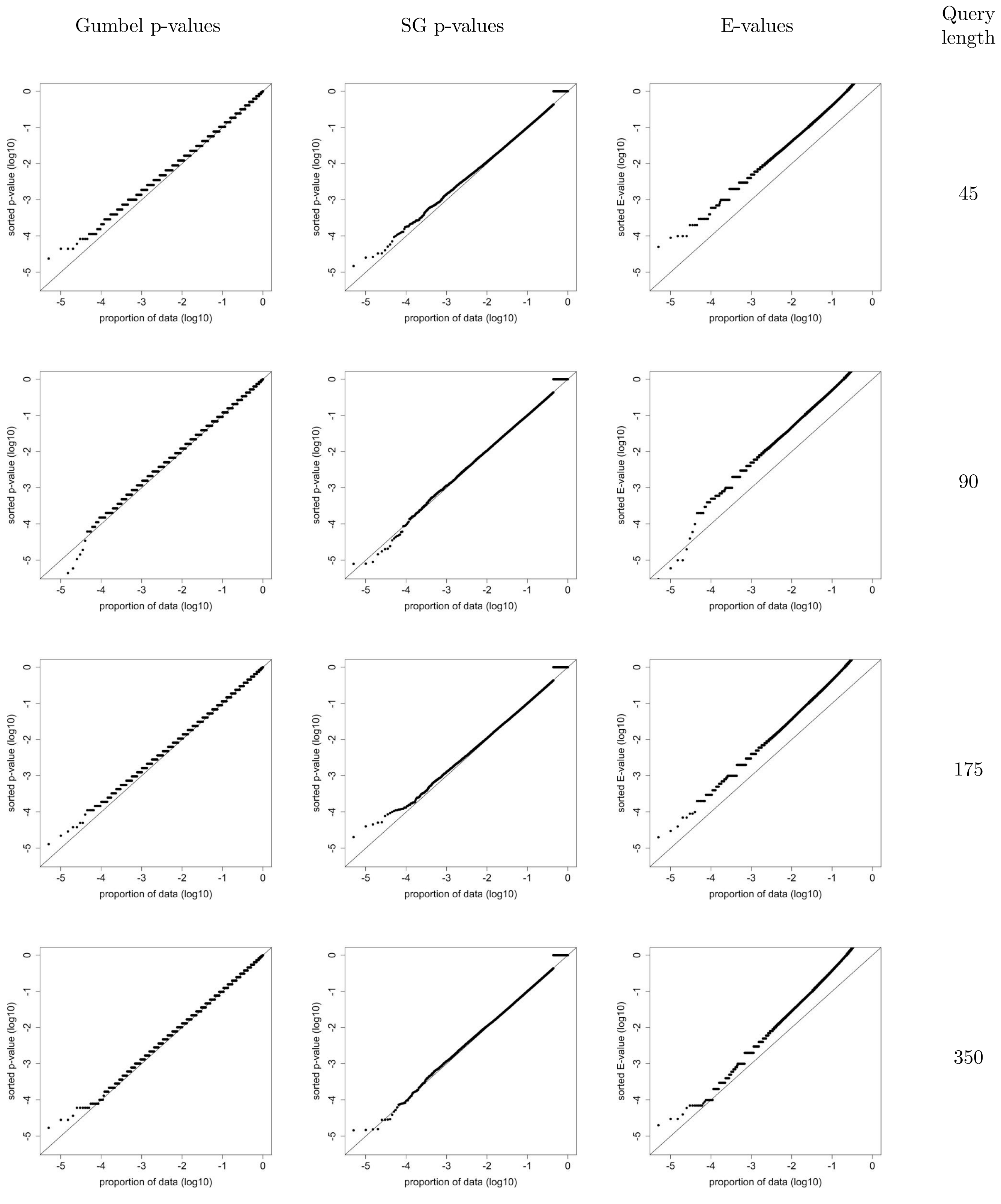


Figure S25: **Using BLAST to search 200,000 shuffled *iid-sampled* queries against the Swiss-Prot database using PAM70 (10,1)**. Same as Figure S7 except using PAM70 (10,1).

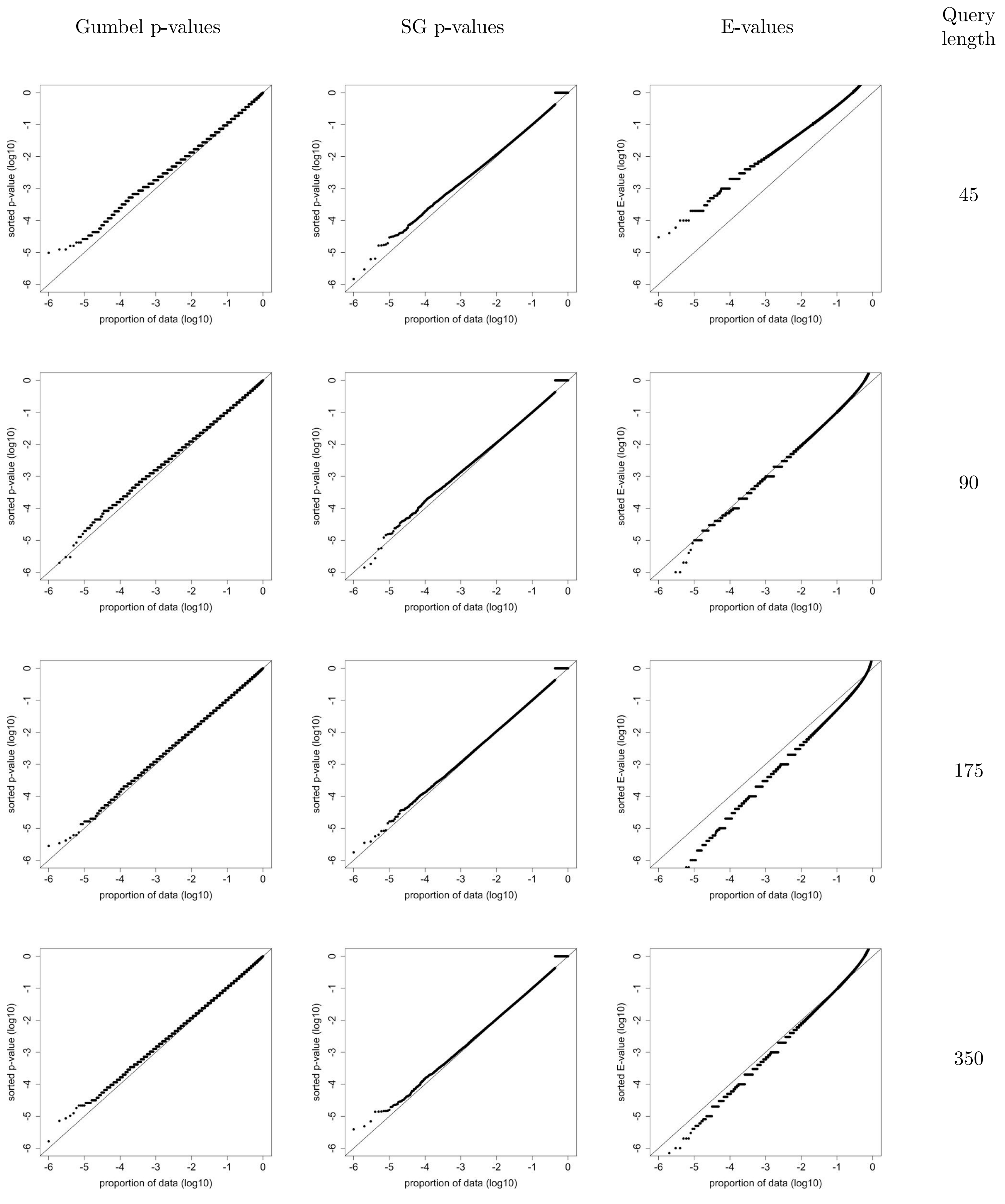


Figure S26: **Using BLAST to search $10^6$ shuffled queries against the Swiss-Prot database with PAM250 (14,2)**. Same as Figure S5 except searching using PAM250. Only 7 of the length 45 shuffles have an E-value $\leq 10^{-4}$ instead of the expected 100. At the same time there are 4.2 times more length 175 shuffles with an E-value $\leq 10^{-3}$ than expected (p-value of 0).

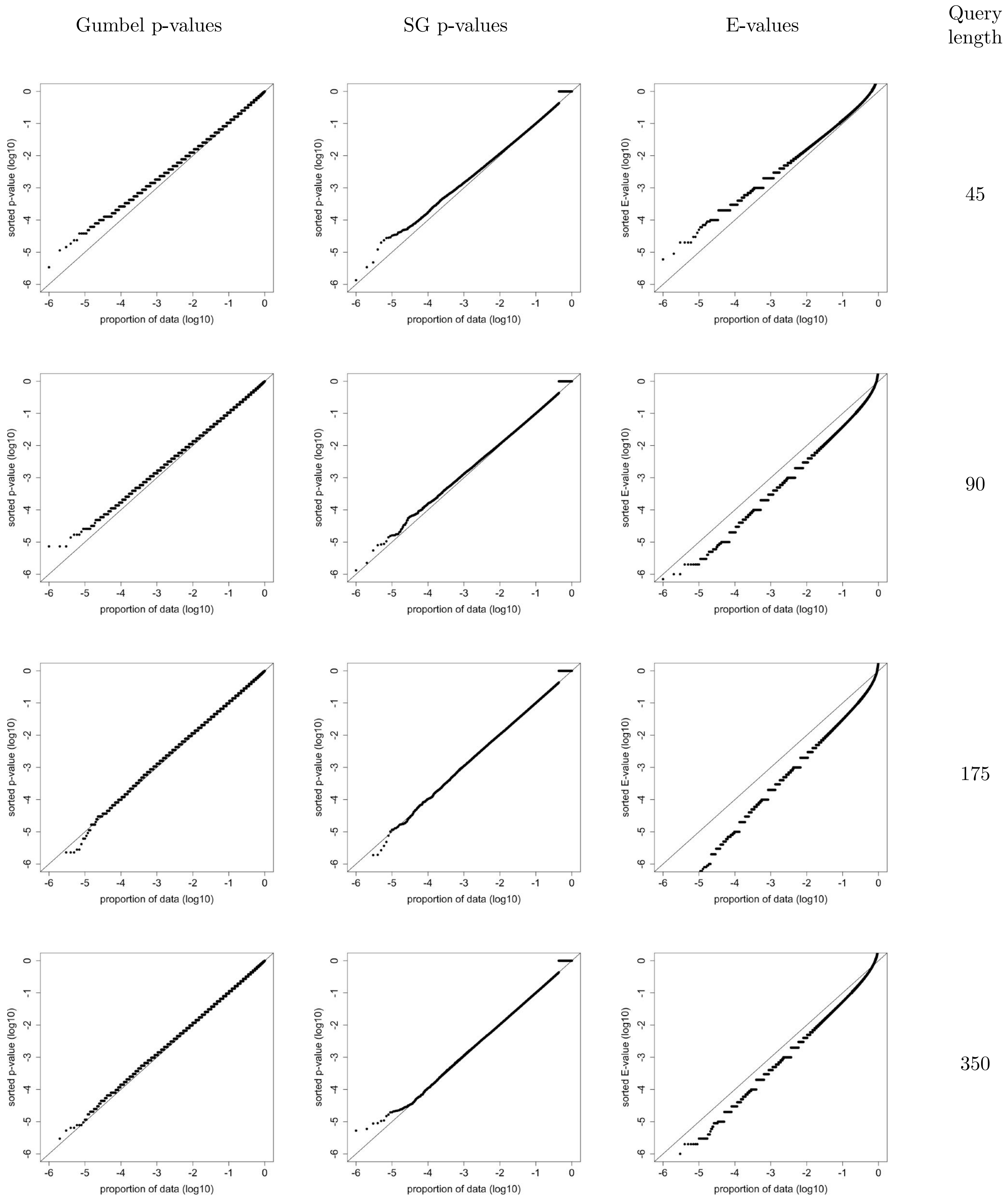


Figure S27: **Using BLAST to search $10^6$ shuffled queries against the SCOP database with PAM250 (14,2)**. Same as Figure S6 except searching using PAM250. There are 15.4% of the length 175 shuffles that with an E-value $\leq 0.05$ (p-value of 0).

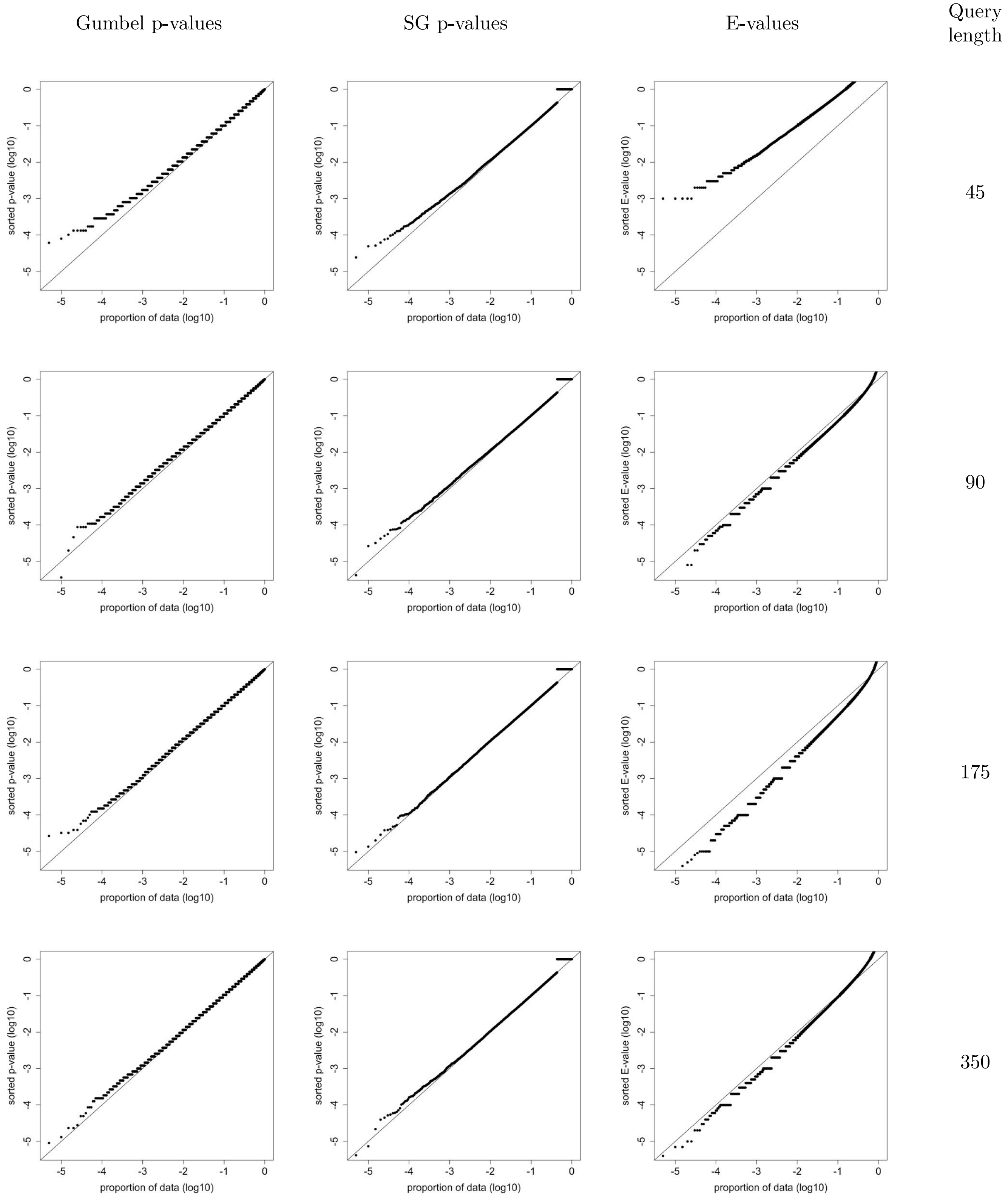


Figure S28: **Using BLAST to search 200,000 shuffled *iid-sampled* queries against the Swiss-Prot database using PAM250 (14,2)**. Same as Figure S7 except using PAM250 (14,2). Only 5 of the length 45 shuffles have an E-value $\leq 10^{-3}$ instead of the expected 200. At the same time there are 4.2 times more length 175 shuffles with an E-value $\leq 10^{-3}$ than expected (p-value of $7 \cdot 10^{-253}$).

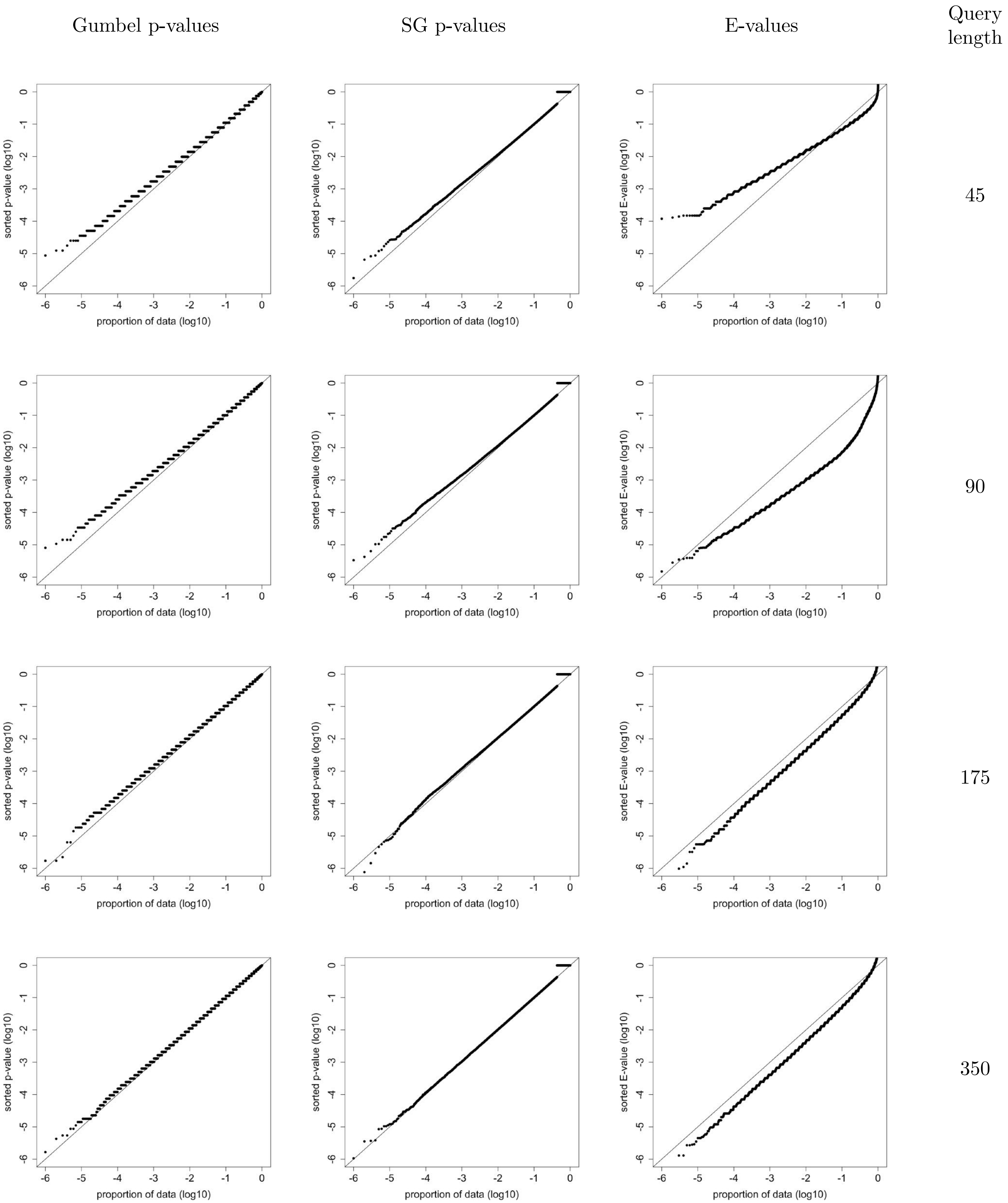


Figure S29: **Using AB-BLAST to search the SCOP database with BLOSUM62 (9,2)**. Similar to figure S6 except the search engine is AB-BLAST using BLOSUM62 with AB-BLAST's default gap penalties of (9,2). Notably AB-BLAST's E-values are not well-calibrated: for queries length of 45 they are at the same time too liberal for some thresholds and overly conservative at others, e.g., 6.1% of the E-values are $\leq 0.05$ (binomial test p-value is 0) but none of the $10^6$ E-values is $\leq 10^{-4}$, and moreover, 13.9% of the queries of length 90 have an E-values $\leq 0.01$ (p-value is 0). The E-values are also clearly too liberal for queries of length 175, e.g., 9.8% of the E-values are $\leq 0.05$ (p-value is 0), and 10% of the E-values of 350-long queries are $\leq 0.05$ (p-value is 0). At the same time the $SG_{50}$ p-values offer a valid and fairly well calibrated alternative for all query lengths.

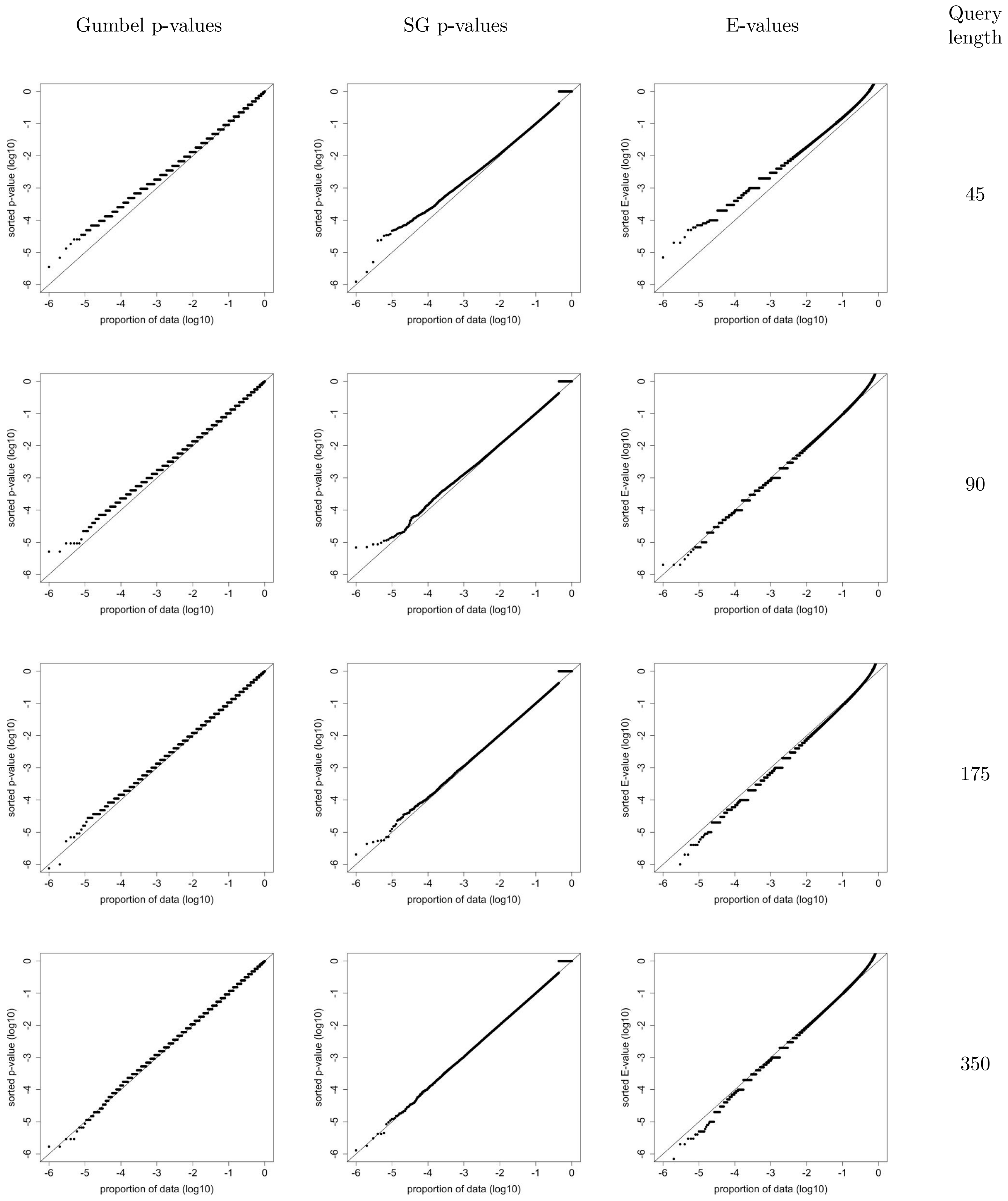


Figure S30: **Using BLAST to search $10^6$ shuffled queries against the ASTRAL40 database with BLOSUM62 (11,1)**. Same as Figure S5 except searching ASTRAL40 instead of Swiss-Prot. Notably the BLAST E-values can be clearly too liberal for the queries of lengths 175, e.g., 6.1% of the E-values are $\leq 0.05$ (binomial test p-value is 0), and length 350, e.g., 5.4% of the E-values are $\leq 0.05$ (p-value is $2.8 \cdot 10^{-87}$). In addition, they are also too liberal for length 90, e.g., 0.18% of the E-values are $\leq 0.001$ (p-value is $5 \cdot 10^{-110}$), and for length 700 (not shown), e.g., 0.13% of the E-values are $\leq 0.001$ (p-value is $1 \cdot 10^{-17}$). In contrast, the $SG_{50}$ p-values are valid and generally better calibrated for all query lengths.

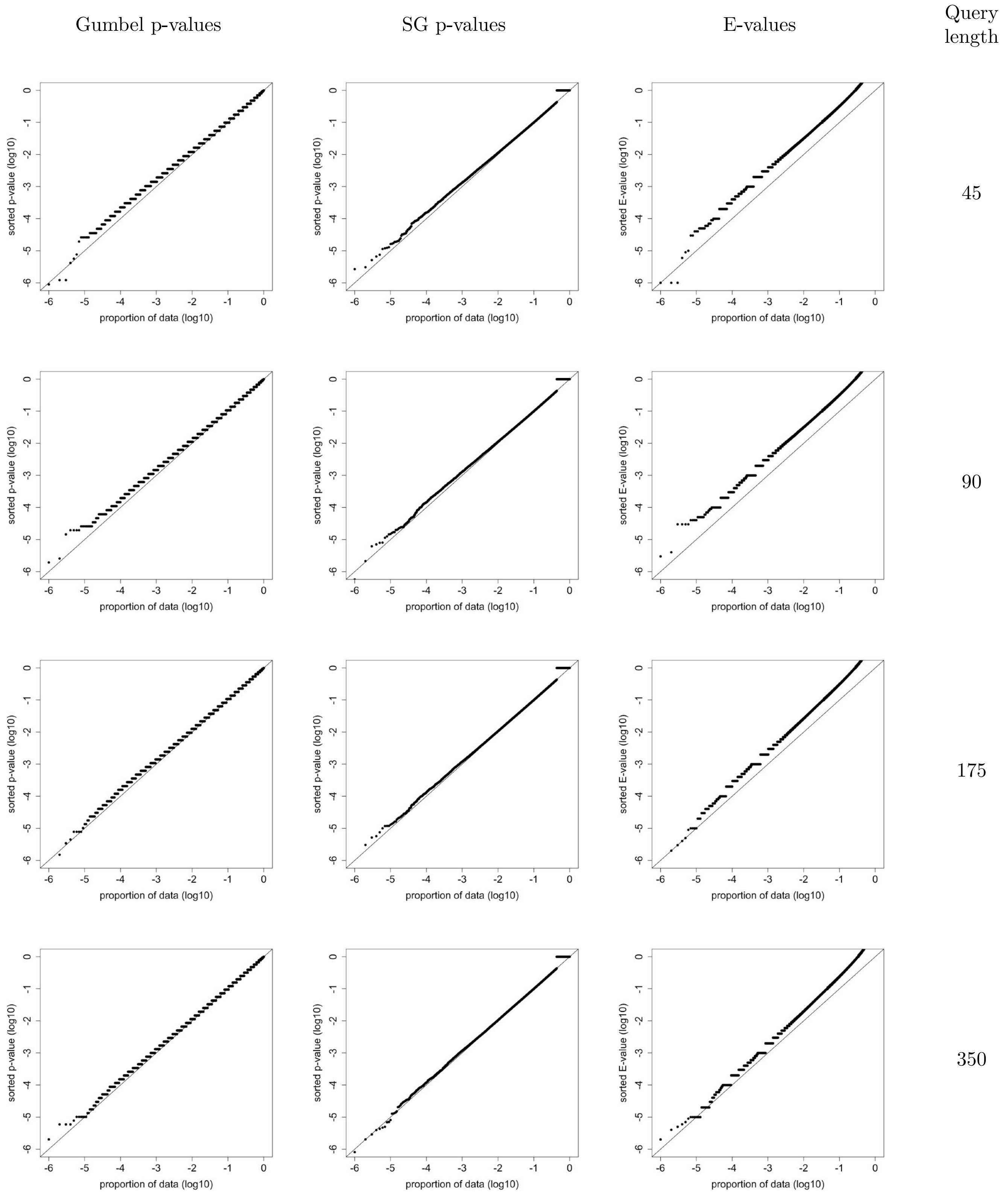


Figure S31: **Using BLAST to search $10^6$ shuffled queries against the ASTRAL40 database with PAM70 (10,1)**. Same as Figure S30 except searching using PAM70 (10,1) instead of BLOSUM62 (11,1). In contrast with the latter case, here BLAST's E-values are consistently conservative. While both the E-values and the $SG_{50}$ p-values are valid here, the E-values are significantly more conservative than the p-values.

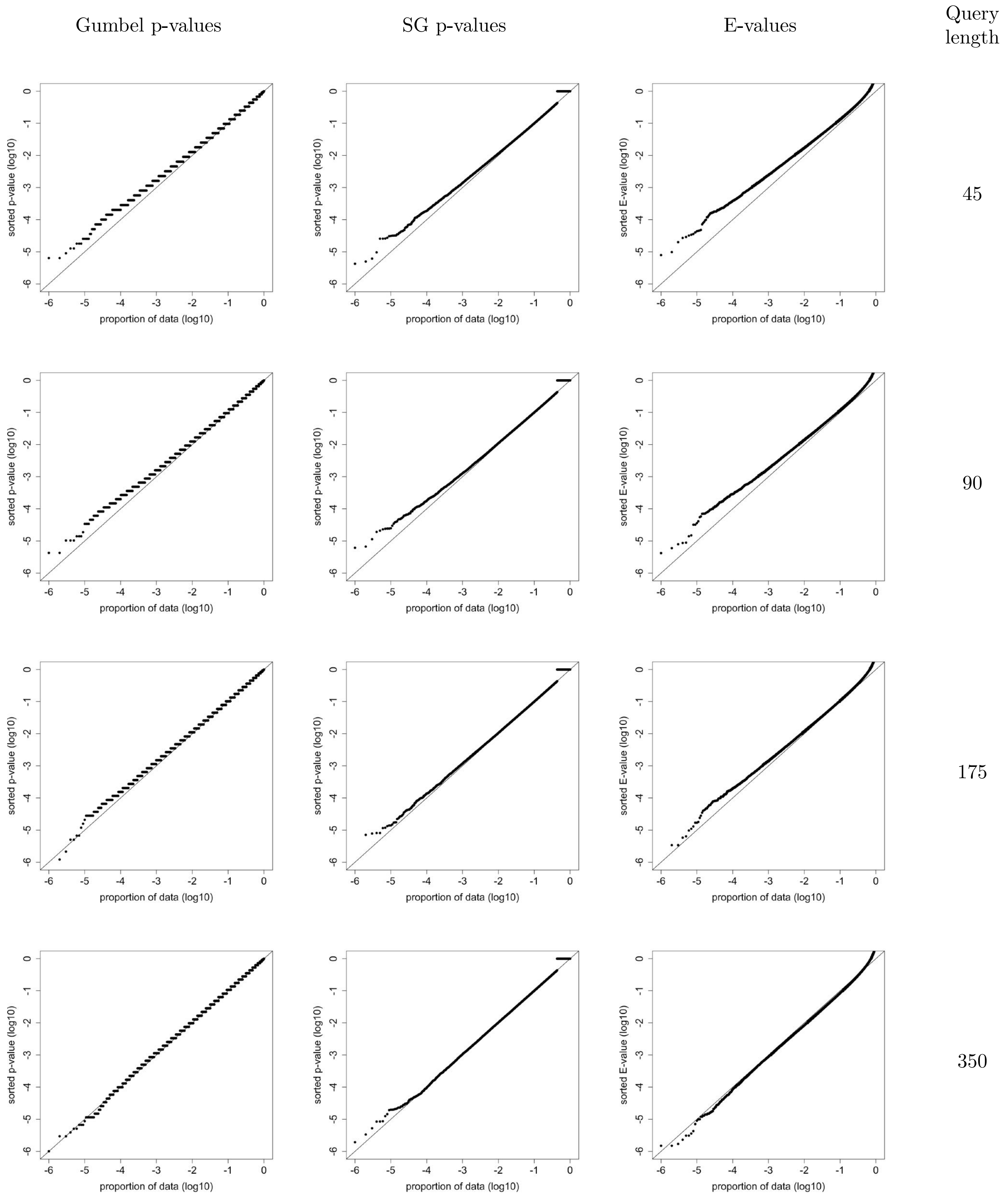


Figure S32: **Using FASTA to search the ASTRAL40 database with BLOSUM62 (11,1)**. Same as Figure S30 except searching with FASTA. While not clearly visible, the FASTA E-values are too liberal for the query of length 350, e.g., 6% of the E-values are $\leq 0.05$ (binomial test p-value is 0), whereas the $SG_{50}$ p-values are (a) valid, and (b) better calibrated for all query lengths.

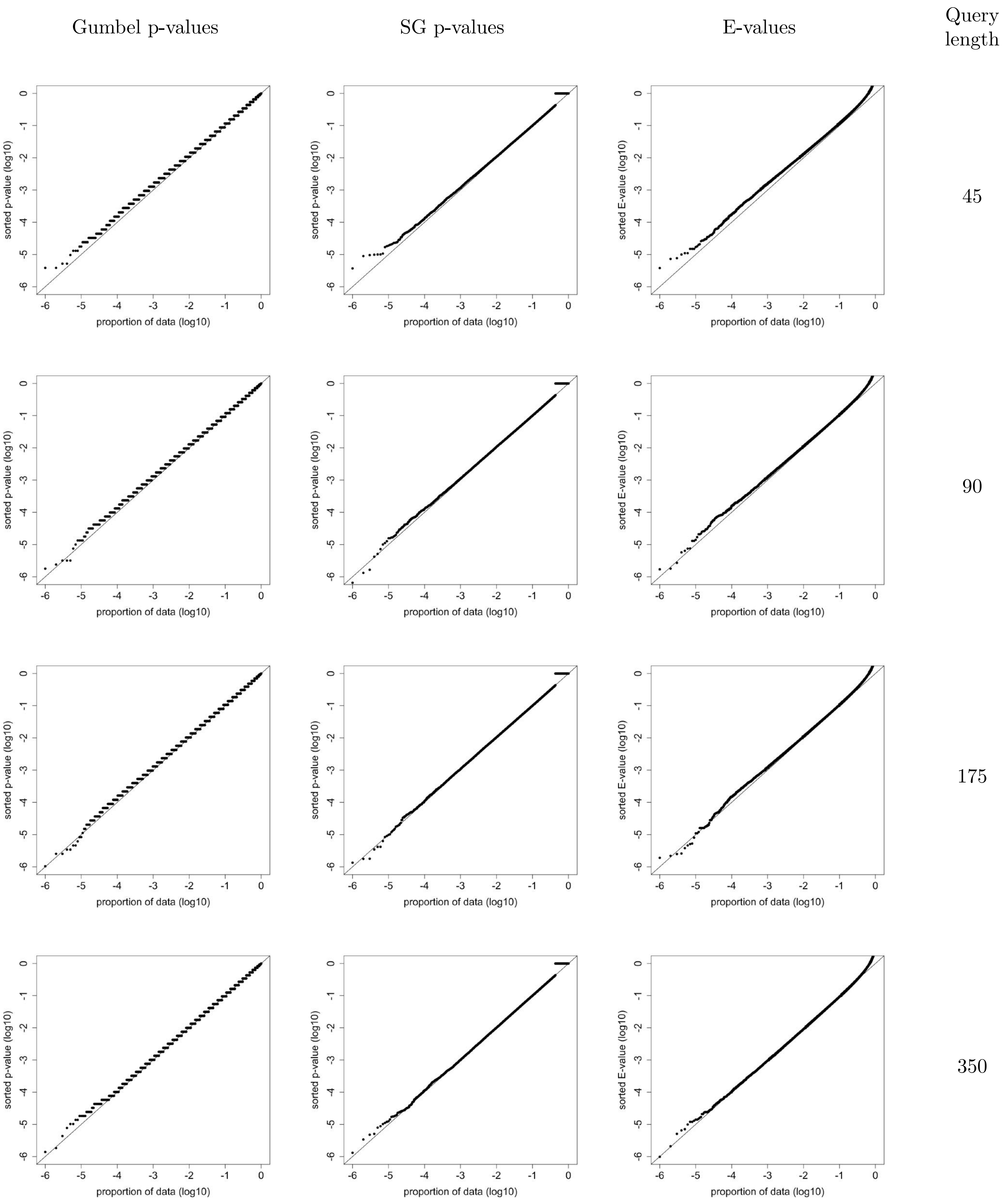


Figure S33: **Using FASTA to search the ASTRAL40 database with PAM70 (10,1)**. Same as Figure S32 except searching with PAM70 (10,1) instead of BLOSUM62 (11,1). Similarly to the latter case, while not clearly visible, the FASTA E-values are too liberal for the query of length 350, e.g., over 5.5% of the E-values are $\leq 0.05$ (binomial test p-value is $1.8 \cdot 10^{-121}$) whereas the $SG_{50}$ p-values are (a) valid, and (b) somewhat better calibrated for all query lengths.

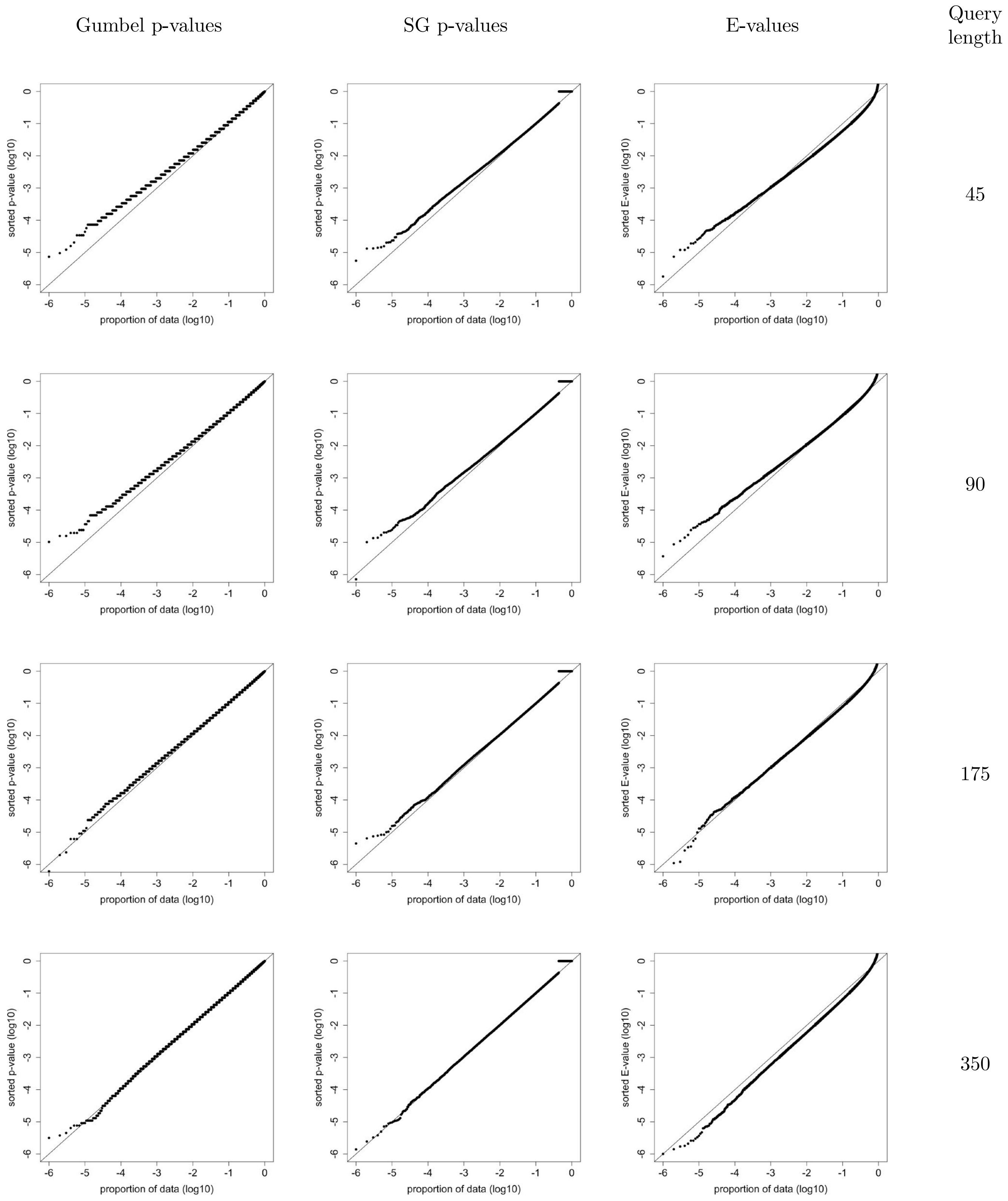


Figure S34: **Using FASTA to search the ASTRAL40 database with PFASUM60 (15,1)**. Same as Figure S32 except searching with PFASUM60 (15,1) instead of BLOSUM62 (11,1). In this case it is clearly visible that FASTA's E-values can be too liberal for length 45, e.g., 8.4% of the E-values are $\leq 0.05$ (binomial test p-value is 0), and length 350, e.g., 8.0% of the E-values are $\leq 0.05$ (p-value is 0). But they are also too liberal for length 90, e.g., 5.6% of the E-values are $\leq 0.05$ (p-value is $1.8 \cdot 10^{-182}$), and length 175, e.g., 6.3% of the E-values are $\leq 0.05$ (p-value is 0). In contrast, the $SG_{50}$ p-values are valid and generally better calibrated for all query lengths.

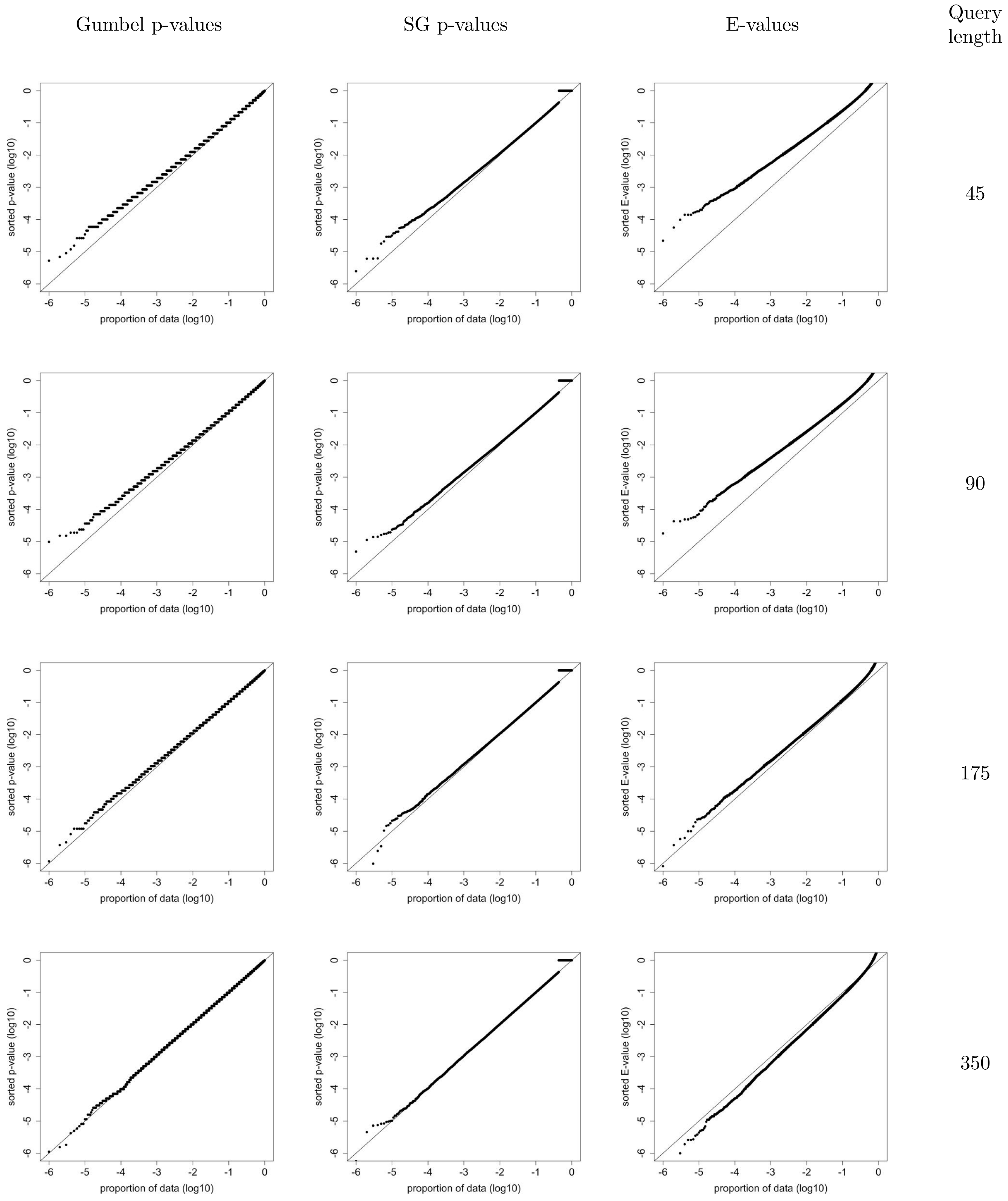


Figure S35: **Using SSEARCH to search the ASTRAL40 database with PFASUM60 (15,1)**. Same as Figure S30 except searching with SSEARCH using PFASUM60 with the (15,1) gap penalties recommended by its authors [15]. Notably the SSEARCH (FASTA) E-values are overly conservative for queries of length 45 they are significantly too liberal for the query of length 350, e.g., 1.5% of the E-values are $\leq 0.01$ (binomial test p-value is 0). In contrast, the $SG_{50}$ p-values are valid and better calibrated for all query lengths.
